# ZmCRY1s interact with GL2 in a blue light dependent manner to regulate epidermal wax composition in *Zea mays*

**DOI:** 10.1101/2025.08.06.668858

**Authors:** Zhiwei Zhao, Fan Feng, Yaqi Liu, Yawen Liu, Fei Wang, Yifan Ni, Huafeng Liang, Wenli Hu, Shanshan Wang, Yuhan Hao, Xu Li, Jigang Li, Jia-wei Wang, Peng Zhang, Hongtao Liu

## Abstract

Cryptochromes (CRYs) are photolyase-like blue-light / ultraviolet-A (UV-A) receptors that regulate diverse aspects of plant growth. Maize (*Zea mays*), a major crop often grown under high UV-B radiation, possesses four copies of CRY. However, it remains unclear whether the multiple copies of CRY in maize have evolved to improve UV tolerance or to acquire new functions. In this study, CRISPR-Cas9-engineered *Zmcry* mutants were used to investigate the functions of four cryptochromes (ZmCRYs) in maize. The findings revealed that ZmCRYs play a redundant role in mediating blue light signaling and in inhibiting the elongation of the mesocotyl. The results also demonstrated that ZmCRYs mediated blue light-enhanced UV-B stress tolerance in *Zea mays* by upregulating the expression of genes involved in UV-B stress tolerance-related metabolites such as phenylpropanoid, flavonoid, and fatty acid biosynthesis. Furthermore, blue light was found to influence both the accumulation and composition of epidermal waxes, suggesting that blue light enhances epidermal wax accumulation for UV-B stress tolerance. Additionally, it was discovered that ZmCRY1 directly interacted with GLOSSY2 (GL2) in a blue light dependent manner to mediate blue light promoted C32 aldehyde accumulation, shedding new light on the enigma of aldehyde-forming. These results highlight the critical roles of ZmCRY1s in mediating blue light regulated epidermal wax biosynthesis and UV-B tolerance in *Zea mays*.

## Introduction

Cryptochromes (CRYs) are photolyase-like blue-light / UV-A receptors that share high sequence and structural similarity with DNA-photolyases ^1–3^. These evolutionarily related flavoproteins perform distinct functions. In Photoreactivation, blue light (350-450 nm) activated DNA photolyase could reversal the harmful effects of far-UV light (200-300 nm), such as growth delay, mutagenesis, and cell death on organisms from bacteria to plants and animals ^3,4^. CRYs regulate growth and development in plants and the circadian clock in plants and animals^5^. In Arabidopsis, two CRY photoreceptors, CRY1 primarily facilitate the plant photomorphogenesis ^6^, while CRY2 regulate the floral initiation ^7–9^. The functionally equivalent dominant mutations in CRY2 boost the adaptation and rapid evolution of species, which may serve as a common genetic basis for ruderality in Brassicaceae ^10^. CRYs are also involved in mediating thermomorphogenesis, and CRY2 protein level is regulated by not only blue light but ambient temperature^11,12^. CRY1 plays role in stomatal opening and mediates various aspects of immunity including pathogen-triggered stomata closure so as to balance photosynthesis and defense^13^. In soil, CRY2 is active in darkness to inhibit root growth demonstrating CRY2’s diverse roles to coordinate developmental processes in above- and below-ground organs ^14^. CRY1 in soybean is essential for blue-light induced root nodule formation, which promotes symbiotic nitrogen fixation in natural ecosystems ^15^. CRY1s regulate the plant height and shade avoidance in soybean and maize^16,17^. Although CRYs lack inherent DNA repair activity, they are nonetheless instrumental in conferring resistance to UV-B damage in plants ^18^.

Maize, a major crop, is grown under the highest light intensity throughout the year and must deal with increased UV-B radiation. The maize genome contains three copies of CRY1 and one copy of CRY2. High-resolution structures of the active PHR domain of ZmCRY1a and ZmCRY1c have been obtained, showing conserved photoactive conformations similar with AtCRYs^19,20^. The conserved photomorphogenesis regulation of ZmCRY1s might be used in breeding of high density-tolerant maize cultivars^16,21^. It is yet to be determined whether multiple copies of CRY in maize have differentiated into new functions or if maize has alternative strategies to cope with high UV-B radiation.

For terrestrial plant, an initial defense against UV-B radiation occurs in the outer cell wall of the epidermis, where less than 10% of incident UV-B is generally transmitted^22^. The epicuticle is composed of various UV-B-screening compounds, including flavonoids, anthocyanin, phenolic acids, and waxes, work complementarily to reflect and absorb UV light ^23–25^. The accumulation or composition of cuticular wax is modulated by varies environmental cues, including different light wavelengths, temperature, osmotic stress^26–31^.various light can enhance wax accumulation by upregulating the transcription of wax biosynthetic enzymes^26,32^. It is still unknown whether CRY could mediate blue light regulated-wax biosynthesis or directly participate the wax biosynthesis.

The chemical composition of waxes primarily consists of very long chain (VLC) aliphatics. Wax biosynthesis begins with de novo fatty acid synthesis of C16 and C18 acyl chains in the stroma of plastids^33^. This process is catalyzed by different types of fatty acid synthase complexes differed in their condensing enzymes which have strict acyl chain length specificities^34,35^: KAS III (C2–C4), KAS I (C4–C16) and KAS II (C16–C18)^36^. The second stage of fatty acid elongation, the extension of the ubiquitous C16 and C18 fatty acids to VLC fatty acids (VLCFAs) chains that are used for the production of aliphatic wax components, is catalyzed through multienzyme fatty acid elongase complexes cycle in endoplasmic reticulum (ER)^33,37^. Multiple β-ketoacyl-CoA synthase (KCS) members have been shown to be involved in fatty acid elongation^38,39^. In the KCS family, KCS5/ ECERIFERUM 60 (CER60) and KCS6/CER6 are major actors involved in the elongation of VLCFAs longer than C26 in Arabidopsis or C28 in maize to produce cuticular waxes with AtCER2/CER2-LIKE / GL2/GL2-like participation^40–43^. VLC acyl-CoAs ranging from 22 to 38 carbon atoms may be processed through two distinct pathways^44^. The alcohol-forming pathway produces even-numbered primary alcohols and alkyl esters, while the alkane-forming pathway produces aldehydes, odd-numbered alkanes, secondary alcohols, and ketones^44,45^. However, the VLC-aldehyde forming enzyme and VLC-alkane synthesis remain largely unknown.

Over time, it has been proposed that alkanes are produced from VLCFAs through the formation of an intermediate aldehydes. CER3 and its homologue CER1 were proved act synergistically as a heterodimer complex with ER-localized cytochrome b5 (CYTB5) as a redox cofactor that catalyzes the presumed decarbonylation of aldehydes to alkanes^46–50^. In addition, two homologs of fatty alcohol oxidase (FAO), FAO3 and FAO4b, are functionally redundant in suppressing accumulation of primary alcohols and contributing to aldehyde production ^51^. Despite these findings, the enzyme responsible for VLC-aldehyde formation and the precise mechanism of VLC-alkane synthesis remain elusive. It has been reported that GL2 overexpression in *cer2-5* mutant or in the Col-0 induces the formation of C32 aldehyde, C31 alkane, C31 secondary alcohol, and C31 ketone ^52^. However, these components were not manifest by the transgenic expression of either AtCER2 or GL2-like. What’s more, the BAHD catalytic HXXXDX-motif is necessary for GL2-like’s function but not for GL2, highlighting the evolutionarily diverged function of GL2 ^52^. Thus, the role of GL2 in alkane-forming pathway remains unclear.

This study reveals the critical roles of ZmCRY1s in mediating blue light enhanced UV-B tolerance in Zea mays. ZmCRYs mediate the blue light-regulated transcription of genes involved in UV-B stress tolerance related metabolites including phenylpropanoid, flavonoid, anthocynin and fatty acid biosynthesis. ZmCRYs mediate the blue light-influenced accumulation and composition of epidermal waxes in maize. Furthermore, ZmCRY1 directly interacts with GL2 in a blue light dependent manner to mediate blue light promoted C32 aldehyde formation, shedding new light on the VLC-aldehyde generation process. Consequently, blue light may play a role in organizing the resistance to UV stress at the outermost layer of the plant, potentially enhancing the plant’s overall tolerance to UV-B radiation.

## Results

### ZmCRYs mediate blue light signaling in maize

To investigate the function of CRYs in maize, we searched maize genome sequence database and classified ZmCRYs family. The maize CRYs are encoded by a multigene family comprising four members and are phylogenetically classified as ZmCRY1s (ZmCRY1a, ZmCRY1b, and ZmCRY1c) and ZmCRY2 (Fig. 1a). ZmCRY1a and ZmCRY1b show 92% sequence identity, they are closely related and likely share similar functions, whereas ZmCRY1c shares 78% and 75% sequence identity to ZmCRY1a and ZmCRY1b respectively. The subcellular localization analysis shows that ZmCRY1a and ZmCRY1b were predominantly localized in the nucleus, with a minor presence in the cytoplasm. In contrast, ZmCRY1c localized in both the nucleus and the cytoplasm, and its proportion in the cytoplasm was significantly higher than that of ZmCRY1a or ZmCRY1b (Supplementary Fig. 1a).

**Fig. 1.**
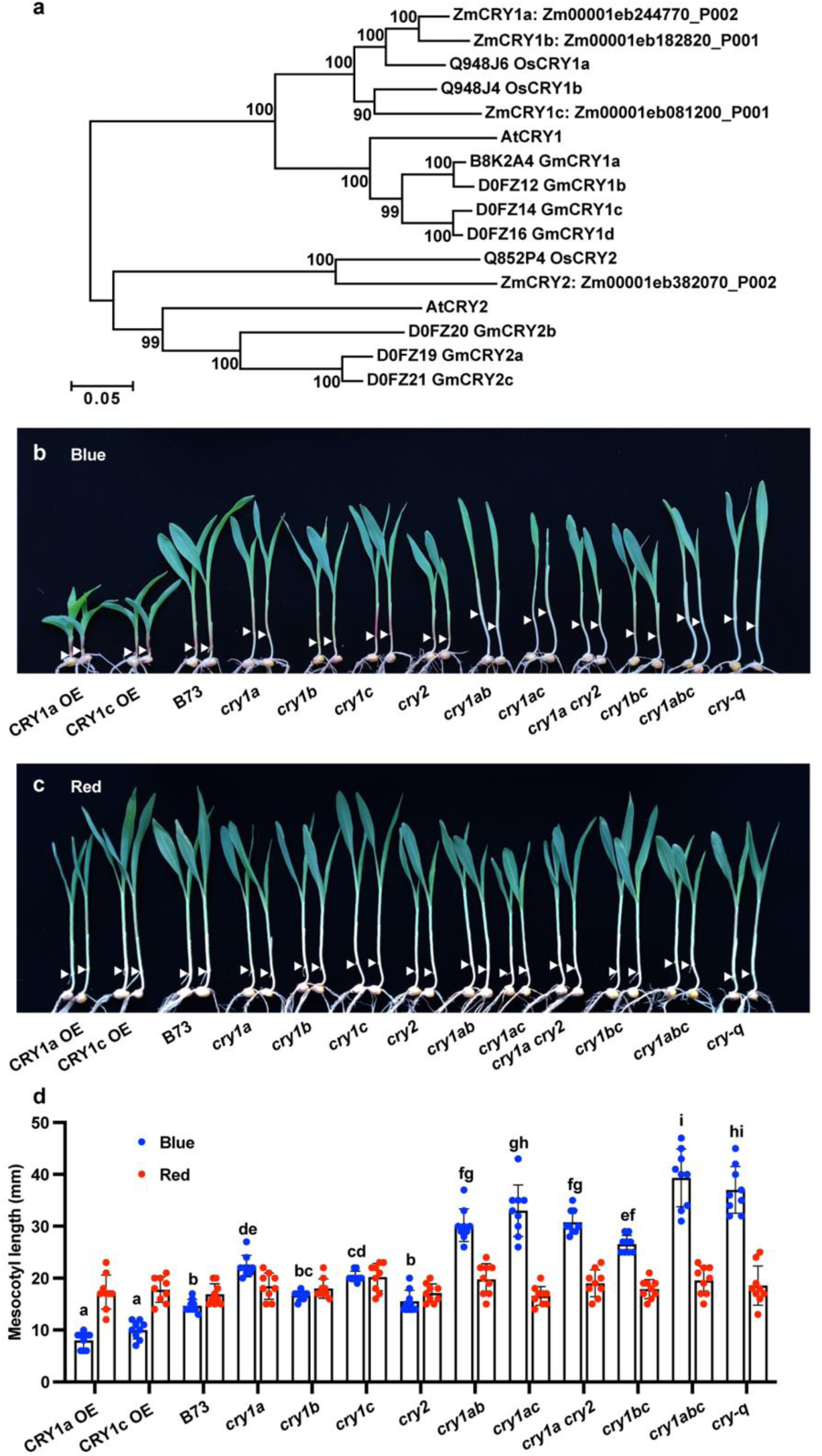
ZmCRYs function redundantly in mediating blue light-repressed mesocotyl elongation in maize. (**a**), Phylogenetic tree of CRY homologues among Maize (*Zea mays*), Rice (*Oryza sativa*), Soybean (*Glycine max (L.) Merr.*) ^65,66^ and Arabidopsis *(Arabidopsis thaliana)*. Distances were estimated using the neighbor-joining algorithm. The numbers at the nodes represent the percentage of 1000 bootstraps. The scale bar indicates the average number of amino acid substitutions per site. **(b and c)**, Representative photographs of indicated genotypes grown under blue light (B, 20 μmol·m^−2^·s^−1^) and red light (C, 20 μmol·m^−2^·s^−1^). **(d)**, Mesocotyl lengths of the indicated genotypes shown in b and c. Error bars represent the s.d. (*n* = 9). Lowercase letters indicate statistically significant differences, no significant differences between genotypes under red light, as determined by one-way ANOVA with Tukey’s multiple comparisons test (*P* < 0.05).

To elucidate their biological functions, uniform Mu insertion lines of CRY1s were obtained from the Maize Genetics Cooperation Stock Center. They were identified that carried Mu insertions at 1377bp (CRY1a: UFMu-08910 named *cry1a-mu*), −10bp (CRY1b: UFMu-09028 named *cry1b-mu*), and 426bp (CRY1c: UFMu-08243 named *cry1c-mu*) from their start codon respectively. PCR, RT–PCR and sanger sequence were performed to examine the existence and the effect of Mu element in CRY1s’ transcripts, no CRY1s’ transcripts were expressed in their Mu insertion lines (Supplementary Fig. 1b to d). Phenotypic analysis revealed that *cry1a-mu* and *cry1c-mu* single mutant showed longer mesocotyl lengths compared to the wild-type W22 inbred line under blue light. The *cry1a cry1c-mu* double mutant show longer mesocotyl length than their single mutants and the wild-type under blue light, while there was no significant difference among the Mu insertion lines under red light (Supplementary Fig. 1e to g). These results suggest that these CRY1s may function redundantly in blue light signaling in maize.

To obtain multiple *cry* knockout mutants, we then employed CRISPR-Cas9 technology to knock out these four cryptochrome genes in B73 background. It was achieved using guide RNA 1 (gRNA1) which targeted conserved regions of ZmCRY1s and gRNA2 which targeted ZmCRY2. *Zmcry quadruple* mutants (*Zmcry-q*) were identified with *ZmCRY1a^d45^*, *ZmCRY1b^d124^*, *ZmCRY1c^d14^* mutants with base deletions and *ZmCRY2^d15,i19^* mutant with 15 base deletion followed with 19 base insertion (Supplementary Fig. 2a). *Zmcry-q* was backcrossed to B73 and self-pollinated to remove the Cas9 transgene and to isolate different single, double, and *Zmcry1s* triple (*Zmcry1abc*) mutants. Compared to wild-type (B73) under blue light in the phenotypic analysis, the single mutants, *Zmcry1a* and *Zmcry1c* showed longer mesocotyl than wild-type in B73 background. Together with double (*Zmcry1ab*, *Zmcry1ac*, *Zmcry1bc*, *Zmcry1acry2*), triple (*Zmcry1abc*), and *quadruple* (*Zmcry-q*) mutants, the mesocotyl length were longer as the functional ZmCRYs decreased. While no significant difference was observed among these genotypes under red light (Fig. 1b to d).

Then we created overexpression lines for *35S::YFP-ZmCRY1a* (CRY1a-OE) and *35S::YFP-ZmCRY1c* (CRY1c-OE) in maize Hi-II background. Three independent transgenic lines were recurrently backcrossed to Chang7-2 for five generations. Phenotypic analysis revealed that the overexpression lines in Chang7-2 background all showed shorter mesocotyl lengths compared to the wild-type specifically under blue light, while no significant difference were observed among these genotypes under red light (Supplementary Fig. 2b to e). To phenotype within the same genetic background, randomly picked CRY1a-OE and CRY1c-OE were recurrently backcrossed to B73 background for five generations. CRY1a-OE and CRY1c-OE showed shorter mesocotyl lengths than B73. These results suggest that ZmCRYs function redundantly in mediating blue light signaling to regulate maize photomorphogenesis.

To analyze whether the blue light receptor function of ZmCRYs is conserved to AtCRYs, we overexpressed ZmCRYs in *Arabidopsis*. YFP-ZmCRY1s mediated blue light inhibition of hypocotyl elongation in transgenic Arabidopsis seedlings, while YFP-ZmCRY2 did not (Supplementary Fig. 3a to c). Photoexcited AtCRY2 proteins become polyubiquitinated and subsequently degraded by the 26S proteasome^53^. AtCRY1 was newly found undergoes degradation in response to high blue light irradiation ^54^. To assess the protein stability of ZmCRYs under blue light, Flag-ZmCRYs overexpression lines in *Arabidopsis* were constructed. The transgenic seedlings were pretreated with 48-hour (h) dark then moved to medium blue light (20 μmol·m^−2^·s^−1^) or high blue light (100 μmol·m^−2^·s^−1^) for 1h time course. When plants were moved to medium blue light, ZmCRY2 protein levels decreased similar as AtCRY2 protein, while ZmCRY1s proteins levels showed no significant decrease (Supplementary Fig. 3 d and e). When plants were moved to high blue light, ZmCRY1a and ZmCRY1b undergoes degradation similar as AtCRY1 protein, while ZmCRY1c was stable (Supplementary Fig. 3 f and g). We suspect the stability of ZmCRY1c may be attributed to its predominantly cytoplasmic localization since the responsible E3 ubiquitin ligases Light-Response Bric-a-Brack/Tramtrack/Broads (LRBs) was nucleus-localized ^12,54,55^. These results suggest that ZmCRY1s and ZmCRY2 are conserved to their blue light receptor orthologous in *Arabidopsis*.

### ZmCRYs mediate the blue light-induced the transcription of genes involved in UV-B stress tolerance in Maize

Current understanding suggests that most CRYs mediated blue light responses in plants are associated with changes in nuclear gene expression. We analyzed the ZmCRYs-dependent blue light-regulated transcriptome. Transcriptome analysis was conducted using 7-day-old long day (LD, 16-hour light/8-hour dark) grown B73 and *Zmcry-q*, pretreated with 48 h dark, then irradiated with or without 1-hour blue light treatment. The transcriptome analysis revealed that blue light regulated a total of 5764 differentially expressed genes (DEGs) in comparison between B73_Blue and B73_Dark (Supplementary Data 1). A Venn diagram analysis indicated that 45.49% (2622) of these DEGs were potentially dependent on ZmCRYs. A heatmap generated from the ZmCRYs-mediated DEGs clearly demonstrated the blue light-regulated genes mediated by ZmCRYs (Fig. 2a, b).

**Fig. 2.**
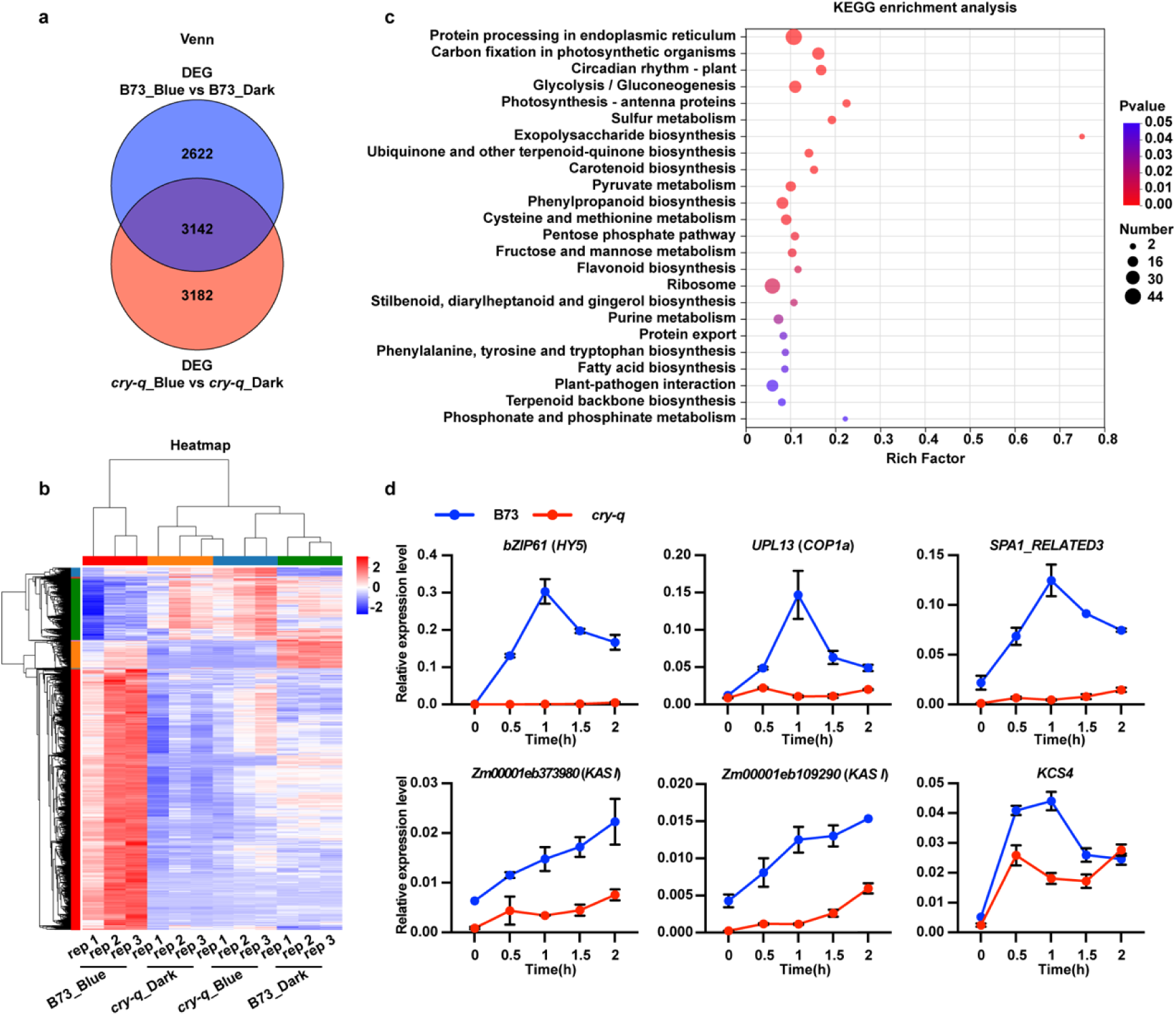
ZmCRYs mediate the blue light-regulated genes’ transcription in Maize. **(a)**, Venn diagram showing the overlap between sets of DEGs from blue light treatment or not in B73 (B73_Blue vs. B73_Dark), and in *cryq* (*cryq*_Blue vs. *cryq*_ Dark) at 28 °C. **(b)**, Heatmap generated from ZmCRYs-mediated DEGs. **(c),** KEGG pathway enrichment analysis of blue up-regulated gene expression. Overrepresented KEGG pathways in the up-regulated gene identified with P value < 0.05 were considered as significantly enriched. **(d)**, RT-qPCR validation the already known blue light signaling pathway regulators and the up-regulated fatty acid biosynthesis enzymes detected in the transcriptome. 7-day LD grown B73 and *Zmcry-q*, pretreated with 48h dark, then moved to blue light (20 μmol·m^−2^·s^−1^) for 2 hours’ time course. *ZmUBQ* served as internal control. Error bars, s.d. of three biological replicates.

Blue light regulates various aspects of plant metabolism, development, and morphology through the regulation of numerous genes’ expression. Addition to the conserved major blue light-responses in *Arabidopsis*, Gene Ontology (GO) enrichment analysis revealed several distinct responses in maize including multiple compounds biosynthesis and metabolic processes (Supplementary Fig. 4 a, b and Supplementary Data 2). It suggests that blue light may promote fatty acid accumulation by enhancing fatty acid biosynthesis while repressing fatty acid beta-oxidation. Additionally, Kyoto Encyclopedia of Genes and Genomes (KEGG) enrichment analysis of blue light-upregulated genes in B73 identified several distinct metabolites related to UV-B stress tolerance, including those involved in phenylpropanoid biosynthesis, flavonoid biosynthesis, and fatty acid biosynthesis as well (Fig. 2c and Supplementary Data 3). These results suggest that fatty acid accumulation may be an important regulation of blue light signaling in maize.

We verified the transcriptome data in seedlings that were moved from dark to blue light (20 μmol·m^−2^·s^−1^) for a 2-hour time course using quantitative reverse transcription PCR (RT-qPCR). For the known blue light signaling pathway regulators detected in the transcriptome, such as the *HY5* homologous gene *BASIC LEUCINE ZIPPER 61* (*bZIP61*) and *bZIP80*, *CONSTITUTIVELY PHOTOMORPHOGENIC 1*(*AtCOP1*) homologous gene *UBIQUITIN*-*PROTEIN LIGASE* 13 (*ZmUPL13*) and *ZmUPL14*, *SUPPRESSOR OF PHYTOCHROME A* 1 (*SPA1*)-*related* 3 and *SPA1-related* 4, *REPRESSOR OF UV-B PHOTOMORPHOGENESIS 2* (*ZmRUP2*) and *CRY3*, their expression was upregulated after blue light irradiation in B73. However, this blue light induction was abolished in *Zmcry-q* mutant (Fig. 2d and Supplementary Fig. 4c). The expression of genes related to fatty acid biosynthesis was also verified using RT-qPCR. *Fatty acyl-acyl carrier protein* (*ACP*) *thioesterase B* (*FatB*), *β-Ketoacyl-ACP-synthase I* (*KAS I*), *β-ketoacyl-ACP-synthase II* (*KAS II*)^34,35^, and *KCSs* (*KCS* genes) were induced by blue light, with partially dependent on CRYs (Fig. 2d and Supplementary Fig. 4d). This suggests that blue light might induce wax accumulation via the regulation of gene expression.

### Blue light affects the accumulation and the composition of epidermal wax in Maize

The expression of genes related to fatty acid biosynthesis was induced by blue light via ZmCRYs, indicating that blue light and CRYs might modulate wax accumulation. To verify this hypothesis, the effects of different wavelengths of light on wax accumulation in maize were analyzed. The maize epidermal wax compositions were measured after 7-day-old dark grown seedlings were exposed to different wavelengths of light (UV-B (1 μmol·m^-2^·s^-1^), blue (20 μmol·m^-2^·s^-1^), red (20 μmol·m^-2^·s^-1^), and far-red light (10 μmol·m^-2^·s^-1^)) for 48 hours. All lights significantly promoted the epidermal wax accumulation compare to darkness (Fig. 3a). The dominant C32 primary alcohol of seedling were significantly elevated in all light treatments. It was observed that blue light and UV-B irradiation had consistent effects on wax composition, with more C32 aldehyde and fewer C24-C26 primary alcohols were accumulated compared to darkness. These results indicate that UV-B, blue, red, and far-red light all promote the accumulation of epidermal wax, while they also show specific effects on the wax composition. To ascertain whether ZmCRYs mediate blue light-regulated wax accumulation, we measured the epidermal wax compositions of 7-day-old seedlings that had been grown in darkness or under continuous blue light, using the CRY1c-OE, B73, and the *Zmcry-q* mutant. There was no significant difference in total waxes content among the different genotypes in either darkness or blue light conditions. This suggests that the blue light-promoted epidermal wax accumulation is not ZmCRYs-dependent (Fig. 3b), and that other blue light receptor families may redundantly regulate wax accumulation. The accumulation of C32 aldehyde were not detected in any genotype under dark. However, under blue light, C32 aldehyde accumulation was much lower in *Zmcry-q* mutant than in B73 and was significantly higher in CRY1c-OE line (CRY1c-OE is 1.45 μg·cm^−2^, B73 is 0.83 μg·cm^−2^, and *Zmcry-q* is 0.33 μg·cm^−2^). These results indicate that the blue light induced C32 aldehyde biosynthesis appears to be primarily ZmCRYs-dependent (Fig. 3b). Furthermore, these findings imply that ZmCRY may mediate blue light responses that participate in the formation of very-long-chain (VLC) aldehydes, a process that remains to be fully elucidated ^44^.

**Fig. 3.**
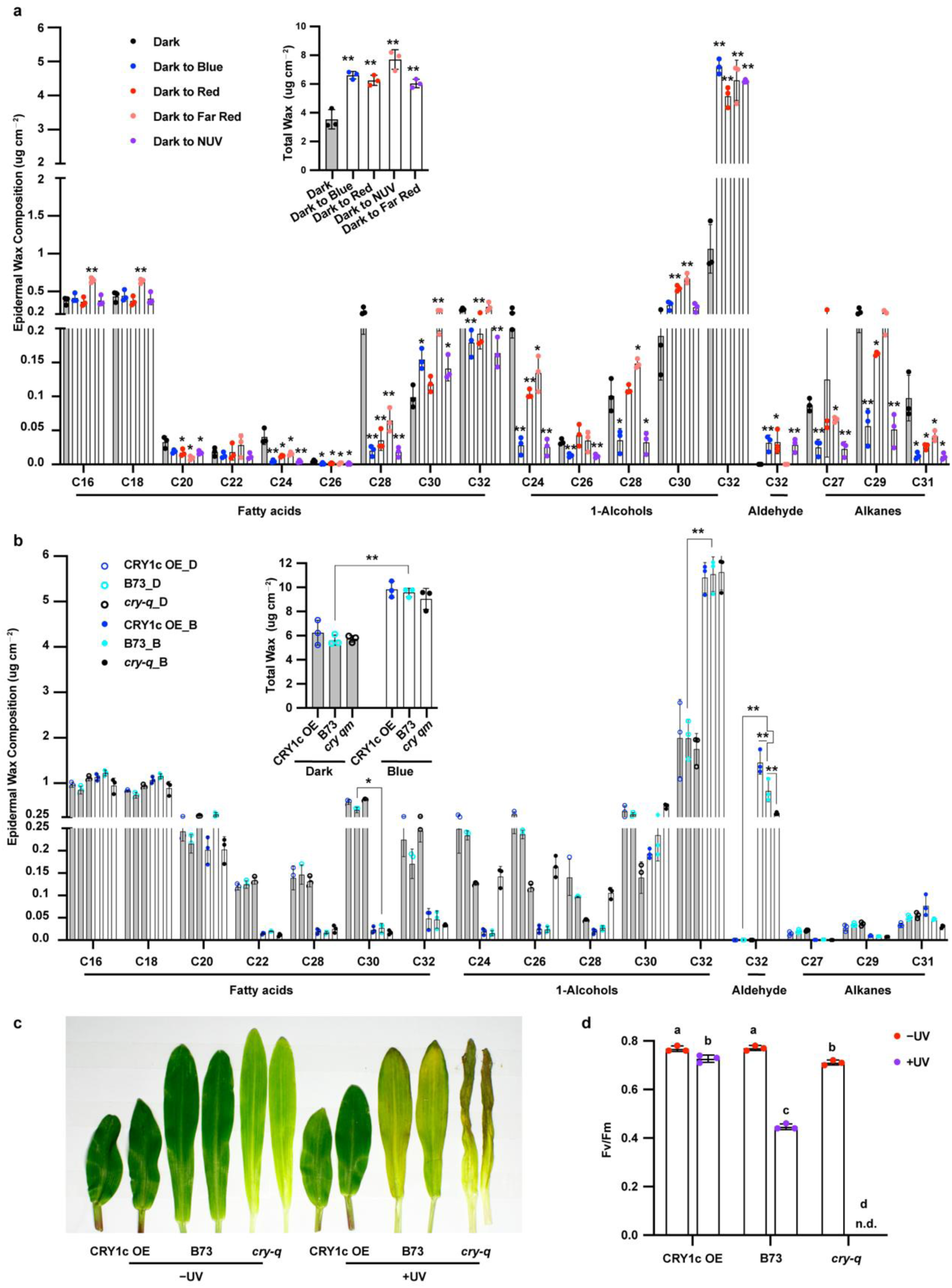
ZmCRYs mediate blue light enhanced epidermal wax biosynthesis. (**a**), Epidermal wax composition analysis of different light. The 7-day-old dark grown seedlings treated with 48h radiation under indicated lights. Blue (Blue light, 20 μmol·m^-2^·s^-1^), Red (Red light, 20 μmol·m^-2^·s^-1^), Far-red (Far-red light, 10 μmol·m^-2^·s^-1^), NUV (Narrow band UV-B light, 1 μmol·m^-2^·s^-1^). Error bars represent standard deviation (s.d., n = 3). The asterisks indicate a significant difference from dark grown B73 based on a two-sided Student’s t-test (*P<0.05, **P<0.01). (**b**), Epidermal wax composition affected by CRY mediated blue light were analysis. 7-day-old of YFP-CRY1c, B73, and *Zmcry-q* were grown in dark or continued blue light (20 μmol·m^-2^·s^-1^). Error bars represent standard deviation (s.d., n = 3). The asterisks indicate statistically significant differences, as determined by two-way ANOVA with Tukey’s multiple comparisons test (*P<0.01, **P<0.001). **(c and d)**, CRY1c enhance Maize tolerance to UV-B stress. **(c)**, Analysis of UV-B stress tolerance in Maize. Plants of the indicated genotypes were grown in soil under long-day (16/8) conditions in blue light (20 μmol·m^−2^·s^−1^) for 10 days and irradiated with (+UV) or without (−UV) broadband UV-B (5 μmol·m^-2^·s^-1^) for 6 hours on day 11 and allowed to recover for 2 days in blue light. **(d)**, Measurement of PSII maximal quantum yield (Fv/Fm) of the first leaves of the indicated genotypes shown in c. Fv/Fm was measured and quantified with an imaging fluorometer. Error bars represent standard deviation. Lowercase letters indicate statistically significant differences, as determined by two-way ANOVA with Tukey’s multiple comparisons test (P < 0.05).

### ZmCRYs enhance the UV-B stress tolerance of maize

Maize is grown under the highest light intensity throughout the year and must cope with increased UV-B radiation. Here, the transcriptome analyzes strengthen that blue light may contribute to UV-B tolerance at the transcription level as reported^18^. To assess the possible UV-B protective capacity of blue light-ZmCRYs, 10-day-old LD blue light (20 μmol·m^−2^·s^−1^) grown CRY1c-OE, B73, and *Zmcry-q* were treated with broadband UV-B light (5 μmol·m^−2^·s^−1^). We measured the maximum quantum yield of Photosystem II (Fv/Fm) as a proxy for UV-B stress tolerance. The results showed that *Zmcry-q* was more sensitive to UV-B stress than B73, while CRY1c-OE exhibited greater tolerance (B73 showed 58.1% resistance from 0.77 to 0.447, CRY1c-OE showed 94.87% resistance from 0.767 to 0.727, and Fv/Fm of *Zmcry-q* were not detected after UV-B irradiation) (Fig.s 3c, d and Supplementary Fig. 5). Additionally, the survival rate of CRY1c-OE was significantly than that of B73 (B73 is 61.9%, CRY1c-OE is 90.5%, and *Zmcry-q* is 33.3% after UV-B irradiation) (Supplementary Fig. 5). In the Chang7-2 background, both CRY1a-OE and CRY1c-OE exhibited enhanced UV-B tolerance compared to Chang7-2. In conclusion, our findings suggest that ZmCRYs mediate blue light-strengthened UV-B tolerance in maize.

Epidermal wax functions as the initial defense against UV-B radiation. Blue light modulates both the accumulation and composition of epidermal wax in maize. However, ZmCRYs primarily mediate the blue light-regulated changes in wax composition, rather than the total wax accumulation. The mechanisms by which ZmCRYs influence waxy composition, as well as the function of C32 aldehyde or its subsequent derivatives in UV-B tolerance, remains to be determined.

### ZmCRY1 physically interacts with Glossy2 in a blue light dependent manner

To investigate how ZmCRY influences waxy composition and the unique functions of ZmCRY1c for the distinct cytoplasmic localization, we employed two strategies to identify the ZmCRY1c-interacting proteins in blue light: yeast two-hybrid screening (Y2H) *in vitro* and *in vivo* co-immunoprecipitation coupled with mass spectrometry (IP–MS). In the Y2H assay, we used ZmCRY1c as the bait to screen a maize cDNA library in blue light (20 μmol·m^−2^·s^−1^). We sequenced 230 positive clones and identified 87 proteins that could interact with CRY1c in blue light in yeast (Fig. 4a, b and Supplementary Data 4). Among these, 4 clones encode various fragments of Glossy2 (GL2), a member of the BAHD acyltransferase superfamily involved in the elongation-reductive pathway of very-long-chain fatty acids. In the IP–MS experiment, we used YFP-ZmCRY1c over-expression line (CRY1c-OE) to affinity purify ZmCRY1c complexes in blue light (20 μmol·m^−2^·s^−1^) or dark. Analysis of the co-purified proteins with CRY1c identified known interacting partners including the COP1 homologue ZmUPL13 and the Gibberellin receptor GID1L2 ^56−58^, as well as the newly identified GL2 (Supplementary Data 4). GL2 specifically enhances the elongation process between chain lengths of C30 and C32 ^59^, and also promotes the formation of C32 aldehyde, C31 alkane, C31 secondary alcohol, and C31 ketone ^52^. Consequently, we selected GL2 as the candidate and conducted further protein-protein interaction experiments to investigate whether blue light promotes the interactions between ZmCRY1c and GL2.

**Fig. 4.**
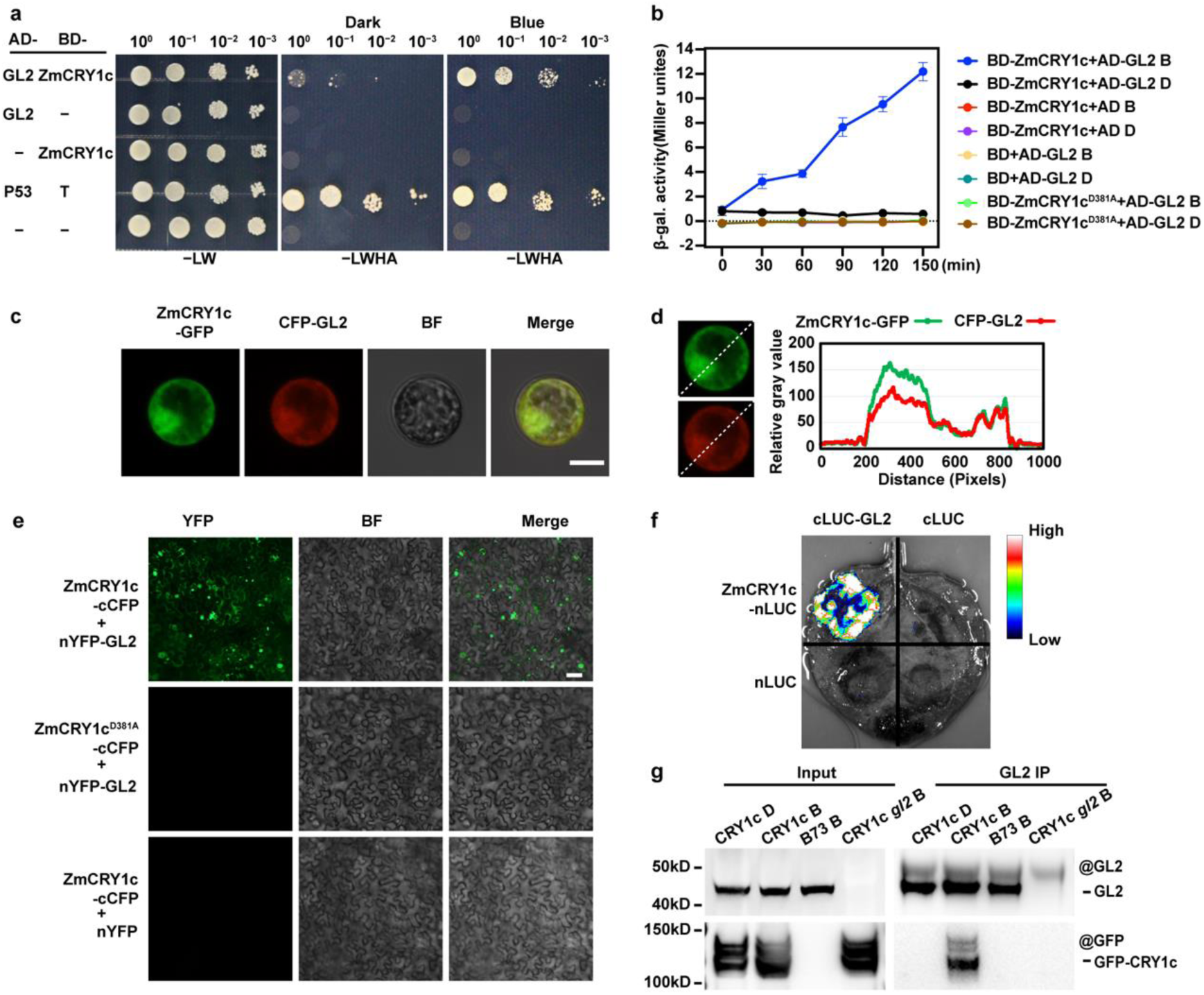
ZmCRY1c interacts with GLOSSY2 in a blue light-dependent manner. **(a)**, Histidine auxotrophy assays showing the interaction between ZmCRY1c and GL2 in a blue light-dependent manner. Yeast cells were grown at –LWHA medium in the dark or under blue light (30 μmol·m^−2^·s^−1^). P53/T was a positive control pair. **(b)**, β-gal assays of yeast cells grown at –LW medium 28 °C in the darkness (D) or under blue light (B, 30 μmol·m^−2^·s^−1^) for the indicated times. Three biological replicates are listed. ZmCRY1c^D381A^ is a site-specific mutant of ZmCRY1c that cannot be activated by blue light. **(c)**, Co-localization of ZmCRY1c and GL2 in cytoplasm in maize protoplast. BF, brightfield. Merge, overlay of the GFP, CFP and brightfield images. Scale bar = 10 μm. **(d)**, Gray-value analysis of fluorescent signals along the dashed diagonal line in the images at left. **(e and f)**, BiLC and BIFC assay showing *in vivo* protein interactions between ZmCRY1c and GL2. Leaf epidermal cells of Nicotiana benthamiana were co-transformed to express the fusion proteins as indicated. In BIFC assay, ZmCRY1c^D381A^-cCFP and nYFP-GL2 group served as negative control. Scale bar = 200 μm. **(g)**, Co-IP assays showing that ZmCRY1c interacts with GL2 in a blue light-dependent manner in plant cells. 7-day-old 28 °C LD-grown seedlings were pretreated in darkness for 48 h, then exposed to blue light (B, 20 μmol·m^−2^·s^−1^) for 2 h. Input: immunoblots showing the abundance of GL2 and YFP-ZmCRY1c in the total protein extracts. GL2 immunoprecipitation (IP): IP products precipitated by the anti-GL2 antibody. Total proteins (Input) or IP products of GL2-beads (GL2 IP) were probed in immunoblots with an anti-GL2 or anti-GFP antibody.

Y2H assays revealed that ZmCRY1c physically interacts with GL2 under blue light, but not in the darkness (Fig. 4a). We quantified the strength of interactions between ZmCRY1c and GL2 by β-galactosidase assay. The interactionss were strengthened only when cells were transferred from darkness to blue light, not while remaining in darkness (Fig. 4b). We failed to detect any interaction between GL2 and the flavin-deficient ZmCRY1c variant ZmCRY1c^D381A^^60^ (Fig. 4b). ZmCRY1c co-localized with GL2 in the cytoplasm of *Nicotiana benthamiana* leaves and maize leaf protoplasts (Fig. 4c, d and Supplementary Fig. 6a, b). Furthermore, we confirmed the interaction between ZmCRY1c and GL2 in plant cells using bimolecular fluorescence complementation (BIFC) assays and bimolecular luminescence complementation (BiLC) assays. Strong yellow fluorescent protein (YFP) fluorescence was observed in the cytoplasm of Nicotiana benthamiana leaves transiently co-infiltrated with ZmCRY1c-cCFP and nYFP-GL2 constructs, indicating a robust interaction, in contrast to leaves co-infiltrated with ZmCRY1c^D381A^-cCFP and nYFP-GL2 (Fig. 4e). Similarly, strong luminescence was observed in leaves coinfiltrated with ZmCRY1c-nLUC and cLUC-GL2 constructs in BiLC assays (Fig. 4f). Additionally, ZmCRY1a and ZmCRY1b were also found to interact with GL2 using BiLC assays (Supplementary Fig. 6c). The *in vivo* interaction of ZmCRY1c and GL2 was confirmed by co-immunoprecipitation assay. Seedlings were pretreated in darkness for 48 hours and then exposed to blue light for 2 hours or kept in darkness before harvesting. Our results showed that ZmCRY1c co-precipitated with GL2 in samples treated with blue light, where as minimal co-precipitated was detected in samples maintained in darkness (Fig. 4g). Collectively, these results reiterate that blue light promotes the interaction between ZmCRY1c and GL2.

The N-terminal PHR domain of ZmCRY1c, which includes the chromophore-binding domain, was sufficient for the interaction between ZmCRY1c and GL2 in yeast cells (Supplementary Fig. 6d), consistent with their blue light dependent interaction. Subsequently, the crystal structure of the constitutive active CRY1c PHR domain (ZmCRY1c-PHR^W368A^ ^20^) in complex with GL2 was determined by Yaqi L., et al. ^60^

### ZmCRY1 modulates aldehyde accumulation together with Glossy2

The interaction interface between ZmCRY1c-PHR^W368A^ and GL2 was identified to primarily involve the α16-α17 loop, α17 of ZmCRY1c (residues 396-419), and α2, β1-β2 loop of GL2 (residues 25-33, 70-76) as described ^60^. To validate this result, we screened for amino acid site mutations in the Maize EMS-induced Mutant Database (MEMD)^61^. Among the 14 EMS mutants of GL2, two mutants, V24M and G106D exhibited a water-beading phenotype similar to that of *gl2* (Fig. 5a, b and Supplementary Fig. 7a). These mutations were found to be in close proximity to Phe71 of GL2, which is the direct interaction site with ZmCRY1c-PHR^W368A^ ^60^ (Supplementary Fig. 7b). To verify the effects of these mutations, we cloned the mutated cDNAs and assessed their impact on the interaction between ZmCRY1c and GL2 using various assays. The GL2^G106D^ mutation severely disrupted the interaction with ZmCRY1c, as evidenced by the inability of the yeast co-transformed with the corresponding constructs group to grow on histidine-deficient medium under blue light in Y2H assays (Fig. 5c). In contrast, the GL2^V24M^ mutation had a milder effect on the interaction with ZmCRY1c compared to the wild-type GL2 (Supplementary Fig. 7c). In BiLC assays, a faint fluorescent signal was detected for ZmCRY1c-nLUC with cLUC-GL2^G106D^ compared to that with cLUC-GL2, indicating a significant reduction in interaction strength. Meanwhile, cLUC-GL2^V24M^ showed a milder decrease in interaction with ZmCRY1c-nLUC (Fig. 5d). Additionally, the influence of these mutations on the interaction with ZmCRY1c-PHR^W368A^ were verified using pull-down analysis. The results showed that the G106D mutation severely disrupted the interaction between ZmCRY1c-PHR^W368A^ and GL2, while V24M mutation had a slightly weaker effect (Fig. 5e). Consistent with these interaction assays, the mutants also displayed deficient epidermal wax compositions comparable to *gl2*, characterized by reduced levels of the major components such as C32 primary alcohol and C32 aldehyde, with accumulation of C22-C30 VLCFAs (Fig. 5f). Collectively, these results underscore the importance of the interaction interfaces (INT) of GL2 with ZmCRY1c for GL2’s function.

**Fig. 5.**
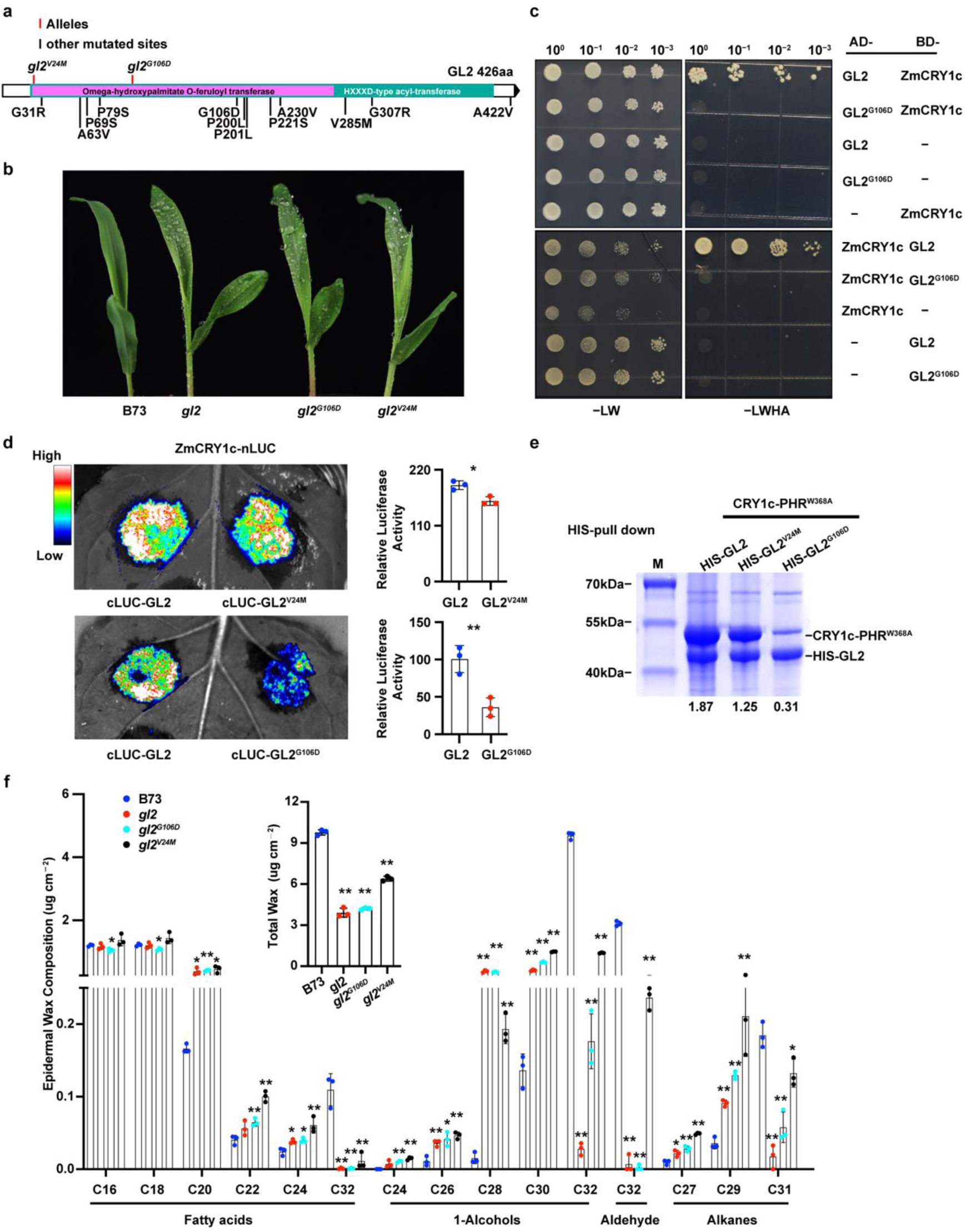
CRY interacting ability is critical for the function of GL2. (**a**), Schematic illustration of GL2 coding sequence with all EMS mutant alleles. **(b)**, Water beading phenotype of the *gl2*, *gl2^G106D^*, and *gl2^V24M^* mutants and wild-type B73 after water spraying. (**c**), Y2H histidine auxotrophy assays showing the site mutation G106D largely disrupts the interaction of GL2 with ZmCRY1c. Yeast cells were grown at –LWHA medium under blue light (30 μmol·m^−2^·s^−1^). **(d)**, BiLC assay showing the site mutation V24M and G106D of GL2 affect its’ interaction with ZmCRY1c in different degrees. A representative picture is show, and the relative luciferase activity are show on the right. Leaf epidermal cells of *Nicotiana benthamiana* were co-transformed with fusion proteins as indicated. Error bars represent the s.d. of three independent biology replicates. The asterisks indicate a significant difference based on a two-sided Student’s t-test (*P<0.05, **P<0.01). **(e)**, HIS-pull down assay on the ZmCRY1c-PHR^W368A^ and GL2 interaction. SDS-PAGE gel shows the influence of His-GL2 site mutations on the complex formation. The numbers below the gels denote the ratio calculated by ZmCRY1c-PHR^W368A^ to His-GL2. **(f)**, Epidermal wax composition analysis of 10-day-old LD white light (50 μmol·m^-2^·s^-1^) grown seedlings of indicated genotypes. Error bars represent standard deviation (s.d., n = 3). The asterisks indicate a significant difference from B73 based on a two-sided Student’s t-test (*P<0.05, **P<0.01).

Overexpression of GL2 in Arabidopsis has been shown to induce the production of C32 aldehyde, C31 alkane, C31 secondary alcohol, and C31 ketone ^52^. To determine whether the accumulation of C32 aldehyde in GL2 overexpression lines is dependent on blue light, we developed GL2 overexpression lines in B73 background. Three overexpression lines were selected for analysis of epidermal wax compositions (Supplementary Fig. 8a). 7-day-old dark or blue light (20 μmol·m^−2^·s^−1^) grown seedlings were sampled, the epidermal wax compositions were measured. The results revealed that the accumulation of C32 aldehyde in GL2 overexpression lines was significantly induced under blue light but abolished in the absence of light, while the induction of C32 1-Alcohol was not dependent on light exposure (Fig. 6a). This indicates that the function of GL2 in promoting the accumulation of C32 aldehyde is regulated by blue light.

**Fig. 6.**
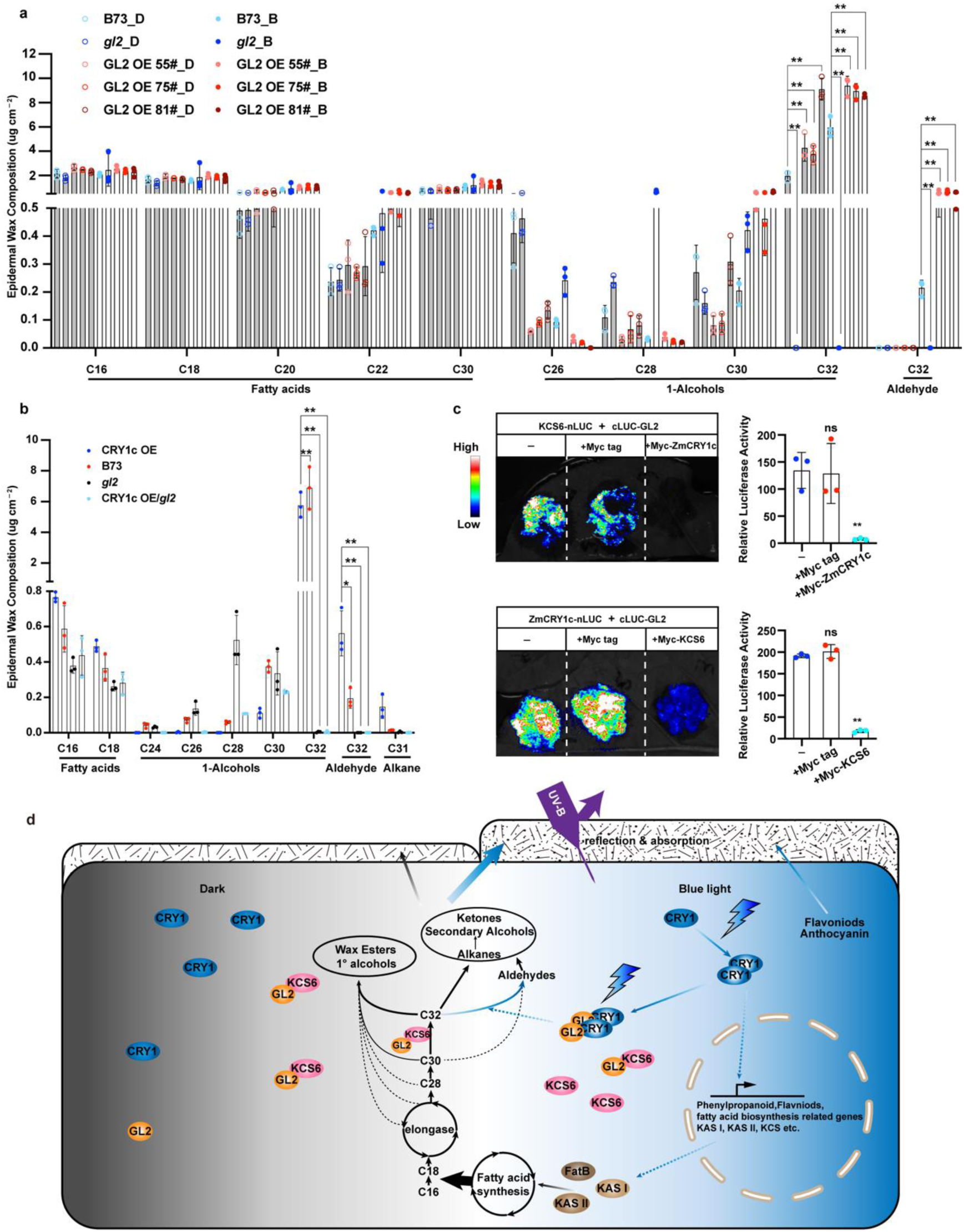
ZmCRY1-GL2 mediate blue light regulated wax biosynthesis. (**a**), Epidermal wax composition analysis of GL2 overexpression lines. D: 7-day-old dark or B: LD blue light grown seedlings of indicated genotypes. Error bars represent standard deviation (s.d., n = 3). The asterisks indicate a significant difference from B73 based on a two-sided Student’s t-test (*P<0.05, **P<0.01). (**b**), Epidermal wax composition analysis of 7-day-old LD blue light (20 μmol·m^-2^·s^-1^) grown seedlings of indicated genotypes. Error bars represent standard deviation (s.d., n = 3). The asterisks indicate statistically significant differences from CRY1c-OE, as determined by two-way ANOVA with Dunnett’s multiple comparisons test (*P<0.01, **P<0.001). (**c**), BiLC assay showing the competitive interaction of ZmCRY1c and KCS6 with GL2. A representative picture is show, and the relative luciferase activity are show below the picture. Leaf epidermal cells of *Nicotiana benthamiana* were co-transformed with fusion proteins as indicated. Error bars represent the s.d. of three independent biology replicates. The asterisks indicate a significant difference based on a two-sided Student’s t-test (*P<0.05, **P<0.01). ns, no statistically significant differences. (**d**), A hypothetical model for blue-light regulation of wax biosynthesis via the ZmCRY1–GL2 signaling pathway.

To determine whether the accumulation of C32 aldehyde in CRY1c-OE is GL2-dependent, we analyzed YFP-ZmCRY1c over-expression in *gl2* mutant backgrounds (CRY1c-OE/*gl2*) (Supplementary Fig. 8b). 7-day-old dark grown seedling were irradiated with blue light (20 μmol·m^−2^·s^−1^) for 48 hours then their epidermal wax compositions were measured. The results revealed that the accumulation of C32 aldehyde in ZmCRY1c over-expression lines was abolished in the absence of GL2, indicating that ZmCRY1c regulates the accumulation of C32 aldehyde in a GL2 dependent manner (Fig. 6b).

### Glossy2 plays dual roles in fatty acid elongation and wax maturation with competitive partners

GL2/CER2-LIKE Proteins play unique biochemical and physiological roles as cofactors with KCS6 in the elongation of VLCFAs up to C34^40−43^. We hypothesized that ZmCRY1c may enhance the interaction between these proteins, thereby facilitating VLCFA elongation. To test this hypothesis, we assessed their interaction using BiLC assays. Unexpectedly, when ZmCRY1c was co-expressed with KCS6-nLUC and cLUC-GL2 in the assay, it diminished the chemiluminescent signal. A similar result was observed when ZmKCS6 was co-expressed with ZmCRY1c-nLUC and cLUC-GL2 (Fig. 6c). These results suggest that ZmCRY1c and KCS6 may compete for interaction with GL2 rather than forming a complex together. Taken together, GL2 interacts with KCS6 to elongate VLCFAs in the dark. Under blue light exposure, the activated ZmCRY1 could interact with GL2 by competing with KCS6, thereby enhancing aldehyde accumulation.

The working model is that blue light activated ZmCRY1 could enhance phenylpropanoid biosynthesis, flavonoid biosynthesis, and fatty acid biosynthesis by promoting the transcription of related enzymes, including *FatB*, *KAS I*, *KAS II* and *KCS*. Additionally, ZmCRY1 can directly interact with GL2 to facilitate the accumulation of C32 aldehyde (Fig. 6d). Thus, blue light induced the epicuticle foundation, contributing to plant tolerance against UV-B stress.

## Discussion

Cryptochromes (CRYs) are blue-light receptors that regulate diverse aspects of plant growth and development, including photomorphogenesis^6^, plant height and shade avoidance^16,17^, floral initiation^7,8,10^. root length and nodule formation^14,15^, among others. This study aimed to characterize the role of CRY in maize. It was discovered that maize CRY mediates blue light-regulated wax biosynthesis and directly participates in the wax biosynthesis process, which may impact important agricultural traits such as UV tolerance, drought stress, and insect resistance.

The RNA-seq data indicates that more compound biosynthesis or metabolic processes were regulated by blue-light in maize compared to *Arabidopsis* (Supplementary Fig. 4a, b and Data 2, 3). The yeast two-hybrid screening and IP-MS results indicate that ZmCRY1c has the potential to interact with numerous proteins and regulate their functions in cytoplasm or membrane system (Supplementary Data 4). This underscores the need for further investigation into the functions of photoreceptors in metabolic processes. The finding expands the effect of light on the agricultural traits. We reported blue light activated ZmCRY1 can influence the accumulation and composition of epidermal waxes in maize. Consequently, blue light may play a crucial role in organizing UV stress resistance at the outermost layer of the plant, potentially enhancing the plant’s overall tolerance to UV-B radiation. We discovered the photoactivated ZmCRY1 interact with GL2, a member of the BAHD family of acyltransferases involved in the elongation of VLCFAs. This ineraction mediates blue light-regulated changes in the composition of epidermal waxes in maize. The study revealed that light can directly regulate fatty acid metabolism via the photoreceptor. However, the ZmCRY1-GL2 interaction appears not conserved at least in *Arabidopsis* ^60^, suggesting that blue light-ZmCRY1-GL2 regulating the composition of epidermal waxes may be specific or evolutionarily diverged in maize.

In addition to various light wavelengths affecting the cuticular waxes in land plants ^29,30^, we further demonstrated that both UV, red, far-red, and blue light all significantly promote epidermal wax biosynthesis and accumulation. Specifically, blue light and UV-B irradiation consistently induce similar changes in wax composition in maize. These observations collectively suggest that different light wavelengths exert both synergistic and specific regulatory effects on the epidermal wax, highlighting the complexity of light-mediated responses in plant cuticles.

A series of wax-deficient mutants have been identified and cloned in *Arabidopsis*, barley, maize, sorghum, and rice, revealing potential functions in wax biosynthesis, transport, or regulation. While the major steps in wax biosynthetic pathway have been characterized ^44,62,63^, the fine-tuning regulation remains to be fully elucidated. The products of alcohol- and alkane-forming pathways form variable wax components depending on environmental factors such as humidity, temperature, or other unknown stimuli. In *Arabidopsis*, the SOH1–CER3–CER1 module has been identfied as a key shunting mechanism. The CER3-CER1 heterodimer catalyzes the decarbonylation of aldehydes to alkanes under water-deficit stress^46−50^, while the CER3-SOH1 complex converts aldehydes into 1-alcohols in response to high-temperature stimuli^27^. These findings emphersize the role aldehydes as central intermediates in the flexible shunting mechanism between alcohol- and alkane-forming pathways, highlighting their importance in plant adaptation to environmental changes. Here we report the mechanism by which blue light promotes the interaction between ZmCRYs and GL2, leading to the induction of C32 aldehyde production thereby enhancing the plant’s adaptation to various environmental stimuli.

KCS6 and GL2 are recognized as major factors involved in the elongation of VLCFA longer than C28 to produce C32 primary alcohols ^40−43^. However, the role of GL2 in alkane-forming pathway remains unclear, as GL2 overexpression induced the formation of C32 aldehyde, C31 alkane, C31 secondary alcohol, and C31 ketone ^52^. In this study, we report that blue light activated ZmCRY1 interacts with GL2 to facilitate the accumulation of C32 aldhyde. Interestingly, ZmCRY1c and KCS6 do not participate in the same complex but instead compete for interaction with GL2. This suggests that GL2 may plays a crucial role in the divergence of VLCFA product pathways, directing them toward the synthesis of alcohols and/or aldehydes in maize. These findings shed new light on the VLC-aldehyde generation process. However, the precise underlying molecular mechanisms remain to be fully elucidated.

## Methods

### Plant materials and growth conditions

Maize (*Zea mays ssp. mays*) inbred line B73, Chang7-2, W22 and hybrid line Hi-II were used. UniformMu insertion lines of CRY1s: UFMu-08910, UFMu-09028, and UFMu-08243 were obtained from the Maize Genetics Cooperation Stock Center. *glossy2* alleles (mutant ID: EMS4-1836a9, EMS4-0eb086, EMS4-02f442, EMS4-0eb085, EMS4-0eb084, EMS4-02f441, EMS4-0eb071, EMS4-0eb070, EMS4-02f427, EMS4-02f426, EMS4-02f425, EMS4-02f423, EMS4-0eb06f) from the MEMD (http://maizeems.qlnu.edu.cn/). *glossy2* was previously described ^59,64^. The transgenic lines and CRISPR-Cas9-engineered mutants were generated in the B73 background.

For UV-B stress analysis in maize, plants were grown at 28 °C, LD conditions in blue light (40 μmol·m^−2^·s^−1^) for 10 d, treated with (+UV) or without (−UV) broadband UV-B (5 μmol·m^−2^·s^−1^) for 6 hours on day 11 and allowed to recover for 2 days in blue light for measurement of PSII maximal quantum yield (Fv/Fm) and allowed to recover for another 2 days prior to phenotypic analysis.

### RNA-seq and Transcriptome analysis

For RNA-seq, B73 and *zmcry-q* seedlings were grown for 7 days LD (white light, 50 μmol·m^−2^·s^−1^) and then pretreated with 48h Dark followed transfer to blue light (20 μmol·m^−2^·s^−1^) for 1 hour. The second leaves were harvested, total RNA was isolated using RNAiso Plus (Takara). Three biological replicates were independently prepared throughout the processes, from the induction of seed germination to the preparation of mRNA-seq libraries. The RNA library generation process followed the manufacturer’s protocol for the Illumina TruseqTM RNA sample prep Kit. DEGs (differential expression genes) with |log2FC|≧1 and FDR < 0.05(DESeq2) or FDR < 0.001(DEGseq) were considered to be significantly different expressed genes. GO functional enrichments were analyzed at https://geneontology.org. KEGG pathway enrichments were analyzed on the online platform of Majorbio Cloud Platform (https://cloud.majorbio.com/).

### Phylogenetic analysis

Amino acid sequences were downloaded from Uniprot (https://www.uniprot.org) and aligned using MUltiple Sequence Comparison by Log-Expectation (MUSCLE) in the MEGA7 software package with the default settings for protein multiple alignment. Evolutionary distances were computed using Poisson correction analysis. The bootstrap method with 1000 replicates for phylogeny testing was used.

### Yeast two-hybrid screening (Y2H)

The Y2H screening assays were performed according to the manufacturer’s instructions (Matchmaker user’s manual, Clontech, California). The sequences encoding CRY1c was fused in-frame with that encoding the GAL4 DNA binding domain (BD) of the bait vector pBridge (Clontech). The resulting constructs was transformed into Y2HGold competent yeast cells. The *Zea mays* whole transcriptome cDNA library cloned in the prey vector pGADT7 was constructed by OE biotech (Shanghai). The resulting constructs were transformed into Y187 competent yeast cells. The Y2H screening were using Mate & Plate™ Libraries according to the manufacturer’s instructions. The screening plates were irradiated with continued blue light (20 μmol·m^−2^·s^−1^). The yeast clones that survived after selection were sequenced, and the correct clones were used in the second round of point-to-point validation.

### Immunoprecipitation–Mass Spectrometry (IP–MS)

2.5 g samples of YFP-ZmCRY1c OE, B73 (7-day old seedlings grown under LD pretreated by 48 h-darkness followed by 2h blue light or maintained in darkness) were collected. The frozen samples were ground into powder in liquid nitrogen and transferred to a mortar with 2 mL extraction buffer A (50 mM Tris pH 7.6, 150 mM NaCl, 5 mM MgCl2, 10% v/v glycerol, 0.5% v/v NP40, 5 mM DTT, 1 mM PMSF, 1× protease cocktail [Roche]). After grinding, the homogenate was incubated on a rotary mixer for 30 minutes 4°C, 35 rpm) and centrifuged at 12,000 rpm for 10 minutes. The supernatant was collected and centrifuged at 12,000 rpm for 10 minutes again. Then, 3mL buffer B (buffer A without NP40) was added, and incubated with GFP agarose beads (25 μL, pre-washed with buffer B for twice) on a rotary mixer for 30 minutes (room temperature, 15 rpm). After incubation, the beads were washed 6 times with washing buffer C (50 mM Tris pH 7.6, 150 mM NaCl, 5 mM MgCl2, 0.2% v/v NP40, 5 mM DTT, 1 mM PMSF, 1×cocktail) and 6 times with buffer D (buffer C without NP40). The resulting beads were eluted by buffer E (2% SDS, 100mM Tris pH 8.0, 10mM TCEP, 50mM CAA) for IP-MS analysis by timsTOF Pro 2 (Bruker).

### BiFC and Subcellular localization assays

In the BiFC assay, constructs for expression of ZmCRY1c or GL2 fused to the C- or N-terminus of YFP were transformed into *Agrobacterium* strain GV3101. For Subcellular localization assays, constructs for expression of ZmCRY1a, ZmCRY1b, ZmCRY1c fused to eCFP were transformed to *Agrobacterium* strain GV3101. For co-localization assay in *Nicotiana benthamiana*, constructs for expression of ZmCRY1c or GL2 fused to CFP or YFP were transformed to *Agrobacterium* strain GV3101. Overnight cultures of agrobacteria were collected by centrifugation, resuspended in MES buffer to 0.6 OD_600_, incubated at room temperature for 2 hours before infiltration. Agrobacteria suspensions in a 1-mL syringe (without the metal needle) were carefully press-infiltrated manually onto healthy leaves of 3-week-old *Nicotiana benthamiana*. Plants were left under LD white light conditions for 2 days after infiltration.

For co-localization assay in Maize protoplast cells, constructs for expression of ZmCRY1c or GL2 fused to GFP or CFP were transformed into *E.coil* MC1601. High concentrations constructs varied between 1 to 2 μg/μL were used for later steps. These constructs were then transformed into protoplasts and culture them for 14-16 hours at 22℃. Finally, collected these protoplasts at 100×g for 2min and removed most supernatant. Observed the YFP signal with confocal microscope (Leica TSC SP8 STED 3X).

### BiLC assays

ZmCRY1a, ZmCRY1b, and ZmCRY1c or GL2, GL2^V24M^, and GL2^G106D^ were fused to sequences encoding the C- or N-terminus of firefly luciferase and transformed into *Agrobacterium* strain GV3101. *Nicotiana benthamiana* plants were left under LD white light conditions for 2 days after infiltration. The leaves were infiltrated with luciferin solution (1 mM luciferin and 0.01% Triton X-100), and images were captured using a CCD camera 5 min later.

### Pull-down assays

The genes encoding ZmCRY1c-PHR^W368A^ with GL2, GL2^V24M^, or GL2^G106D^ were cloned into pET-duet vector, with a 6×His tag at the N terminus of GL2s. E. coli BL21 (DE3) strain was used for protein expression. The protein expression and purification was carried out as described^60^.

### Immunoblot

Immunoblot was carried out as described previously. For anti-ZmCRY1c-CCE and anti-GL2 antibody production, the coding sequence of ZmCRY1c-CCE and full length GL2 were cloned into pET28a (Novagen). Antibody production were performed by ORIZYMES (Shanghai). The anti-ZmCRY1c-CCE and anti-GL2 antibody was used at 1:3000. The anti-GFP (Abicode, #M0802-3a) and anti-Flag (Smart-Lifescience, #SLAB01C) antibody were used at 1:3000.

### Analysis of wax composition

Wax extraction and gas chromatography-mass spectrometry (GC-MS) analyses were performed according to the described methods with some modifications ^64^. In different assays, the horizontal expansion part of the second leaves (2-4 cm from tip) were taken and 4 leaves were collected as one replicate and immediately immersed in 10 ml chloroform with 20 μg of N-tetracosane (CATO) as an internal standard for 30-60 s. The extracts are volatilized drying in a fume hood, and transfer to a 2 mL tube. Dissolved in 60 μL of pyridine and derivatized by adding 60 μL of N, N-bis-trimethylsilyltriflu oroacetamide (Macherey-Nagel), incubated at 70 °C for 1 hour. These derivatized samples were filtered and analyzed by GC with a flame ionization detector (Agilent GC 7890, Technologies) and GC-MS (Agilent gas chromatograph coupled to an Agilent MSD 5977B quadrupole mass selective detector).

### Accession Numbers

Sequence data from this work can be found in the ensembl.gramene or Uniprot under the following accession numbers: Zm00001eb244770 (ZmCRY1a), Zm00001eb182820 (ZmCRY1b), Zm00001eb081200 (ZmCRY1c), Zm00001eb382070 (ZmCRY2), Zm00001eb071110 (GLOSSY2), Zm00001eb008920 (ZmKCS6), AT4G08920 (AtCRY1), AT1G04400 (AtCRY2), GLYMA_04G101500 (GmCRY1a), GLYMA_06G103200 (GmCRY1b), GLYMA_14G174200 (GmCRY1c), GLYMA_13G089200 (GmCRY1d), GLYMA_10G180600 (GmCRY2a), GLYMA_02G005700 (GmCRY2b), GLYMA_20G209900 (GmCRY2c), Os02g0573200 (OsCRY1a), Os04t0452100 (OsCRY1b), Os02g0625000 (OsCRY2). RT-qPCR: Zm00001d015743 (bZIP61) and Zm00001eb385610 (bZIP80), Zm00001d018207 (ZmUPL13) and Zm00001d052138 (ZmUPL14), Zm00001d014990 (ZmRUP2), Zm00001eb361760 (SPA1-related 3), Zm00001eb154420 (SPA1-related 4), Zm00001eb389640 (CRY3), Zm00001eb278580 (KAS I), FABFs Zm00001eb373980 (KAS II), Zm00001eb278580 (KAS II), Zm00001eb364580 (Stearoyl-acyl-carrier-protein desaturase 2, SAD, FAB2), Zm00001eb277000 (FATB), Zm00001eb377350 (FATB), Zm00001eb296230 (KCS4), Zm00001eb344070 (KCS).

RNA-seq data are available from National Center for Biotechnology Information Gene Expression Omnibus (http://www.ncbi.nlm.nih.gov/geo) under the series entries GSE285557.

### Statistical analysis

For phenotype analysis, gene-expression analysis, and statistical analysis were assessed as described in the Fig. legends. *P* values were calculated by two-sided Student’s t-tests or by one-way ANOVA with Tukey’s multiple comparisons test using GraphPad Prism9 and were shown in bar graphs or source data.

## Acknowledgements

The authors thank Dr. Jun Zheng in Chinese Academy of Agricultural Sciences (CAAS) for the *glossy 2* mutant line, thank Dr. Chunyi Zhang in CAAS and Dr. Xiaoduo Lu in QILU normal university for the *glossy* 2 EMS alleles. This work was supported in part by the National Key R&D Program of China (2024YFA1306700 to H.L), National Natural Science Foundation of China (32330006, 31825004, 32150007, 32100197, 32300284, 32170248). Z.Z is supported by the foundation of Youth Innovation Promotion Association of CAS.

## Author contributions

Z.Z., F.F., and H.L. conceived the project. Z.Z., F.F., and Y.w.L. performed most of the experiments, Y.q.L. performed the pull-down assay and Structure analysis. Y.q.L., Y.w.L., Y.N., H.f.L. and Y.H. provided materials. W.H., S.W., X.L., J.L., and J.W. provided technical assistance. Z.Z., P.Z., and H.L. analyzed the data. Z.Z, and H.L. wrote the manuscript.

## Additional information

Correspondence and requests for materials should be addressed to H.L.

