## Supplementary_Materials for "ZmCRY1s interact with GL2 in a blue light dependent manner to regulate epidermal wax composition in *Zea mays*"

**This PDF file includes:**

Supplementary Fig. 1 to 8

**Other Supplementary Materials for this manuscript include the following:**

Supplementary Table 1 to 6 (.xlsx)

Supplementary Table 1. List of differentially expressed genes

Supplementary Table 2. GO enrichment in the differentially expressed genes

Supplementary Table 3. KEGG pathways enrichment in the differentially expressed genes

Supplementary Table 4. List of ZmCRY1c interacting proteins

Supplementary Table 5. List of potential CRY1c-interacting proteins identified by IP–MS

Supplementary Table 6. Primers used in this study.

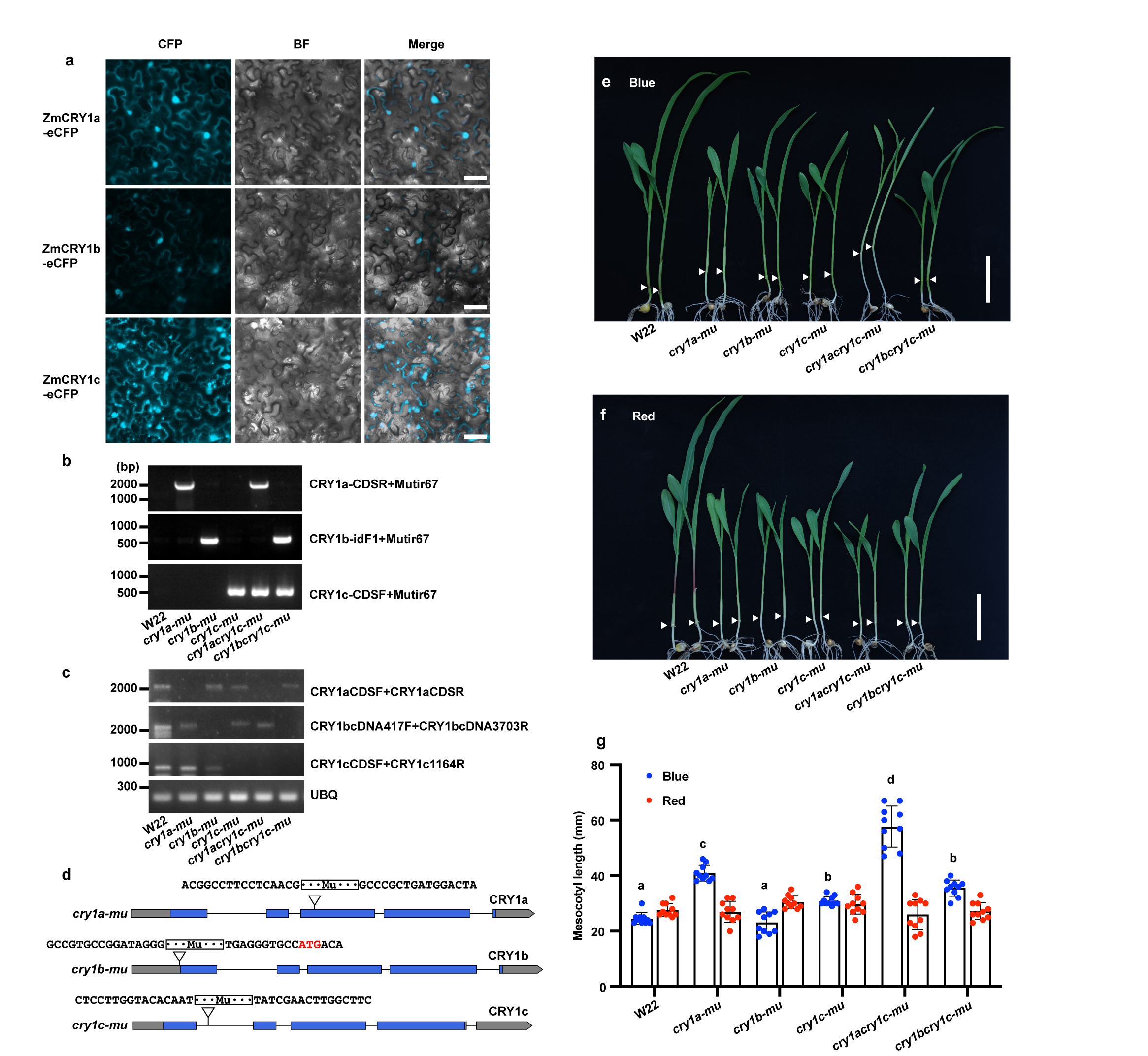

Supplementary Fig. 1 Subcellular localization of ZmCRY1s and Phenotypic analyses of maize *cry1s-mu* mutants. related to Fig. 1.

**(a)**, The subcellular localization of ZmCRY1a, ZmCRY1b and ZmCRY1c in *N. benthamiana* leaves. BF, brightfield. Merge, overlay of the CFP and brightfield images. Scale bar = 50 μm. **(b)**, PCR results showing the genotyping of the indicated mutant. **(c)**, RT-PCR results showing mRNA expression of *CRY1s* in the indicated genotypes. **(d)**, A diagram illustrating the genomic structure of CRY1s and the locations of the Mu insertions. **(e and f)**, Representative photographs of indicated genotypes grown under blue light (e, 20 μmol·m^−2^·s^−1^) or red light (f, 20 μmol·m^−2^·s^−1^) for 7-day. White arrow heads indicate boundary between mesocotyl and first internode. Scale bars, 5 cm. **(g)**, Mesocotyl lengths of the indicated genotypes shown in E and F. Error bars represent the s.d. (*n* = 10). Lowercase letters indicate statistically significant differences, no significant differences between genotypes under red light, as determined by two-way ANOVA with Šídák's multiple comparisons test (*P* < 0.05).

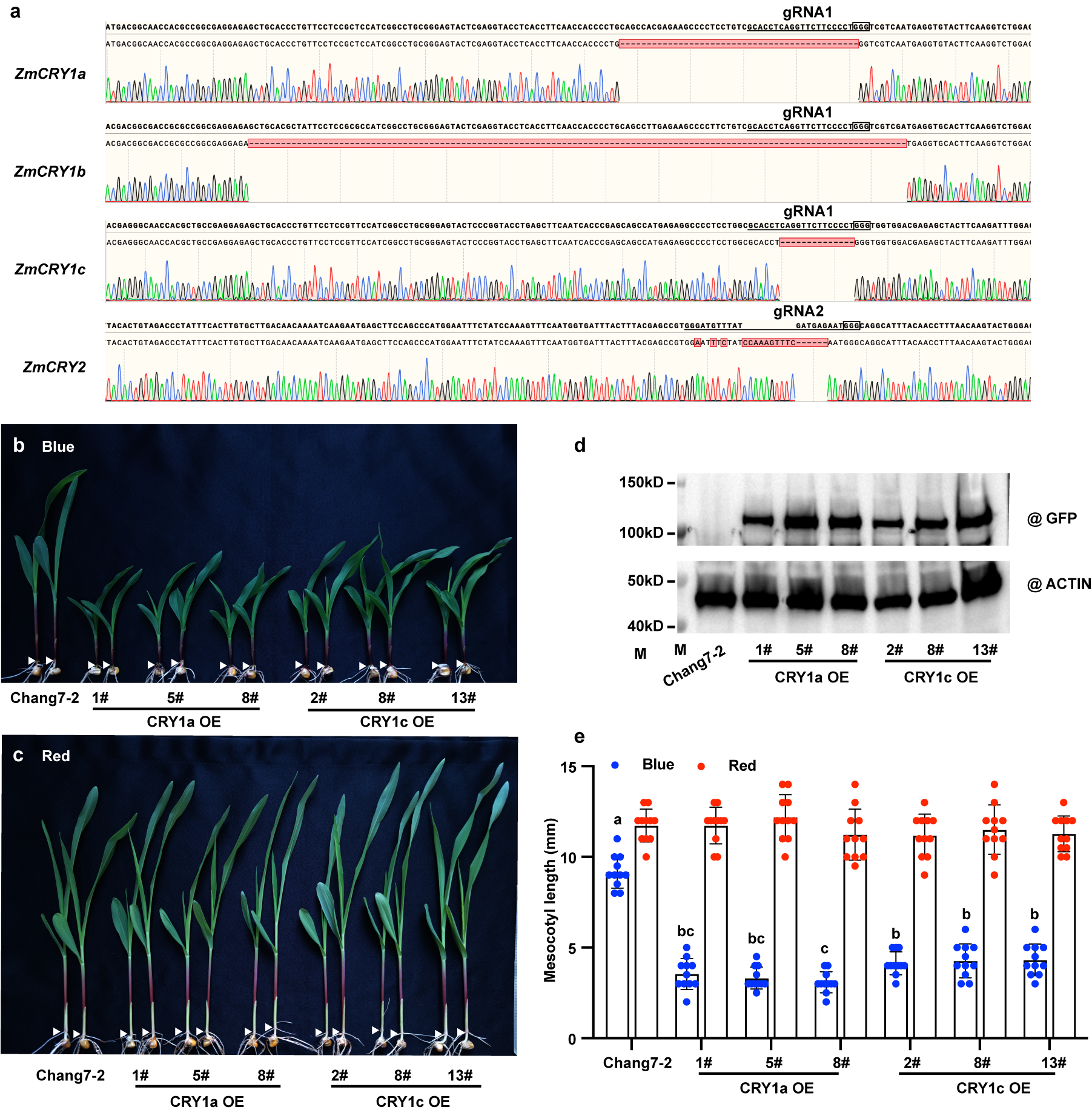

Supplementary Fig. 2 ZmCRY1s mediate blue light-repressed mesocotyl elongation. related to Fig. 1.

(**a**), Knockout mutants of CRYs edited by CRISPR-Cas9 in maize. Sanger sequencing chromatograms of CRISPR-Cas9 edited ZmCRYs genes. guide RNA 1 (gRNA1) which targeted conserved regions of ZmCRY1s and gRNA2 which targeted ZmCRY2 sites are marked. **(b and c)**, Representative photographs of indicated genotypes in Chang7-2 background grown under blue light (20 μmol·m^−2^·s^−1^) and red light (20 μmol·m^−2^·s^−1^). **(d)**, Immunoblots showing the YFP-ZmCRY1a and YFP-ZmCRY1c protein level in their overexpression transgenic lines in Chang7-2 background as indicated. ACTIN serves as the loading control. **(e)**, Mesocotyl lengths of the indicated genotypes shown in (b and c). Error bars represent the s.d. (*n* = 11). Lowercase letters indicate statistically significant differences, no significant differences between genotypes under red light, as determined by one-way ANOVA with Tukey’s multiple comparisons test (*P* < 0.05).

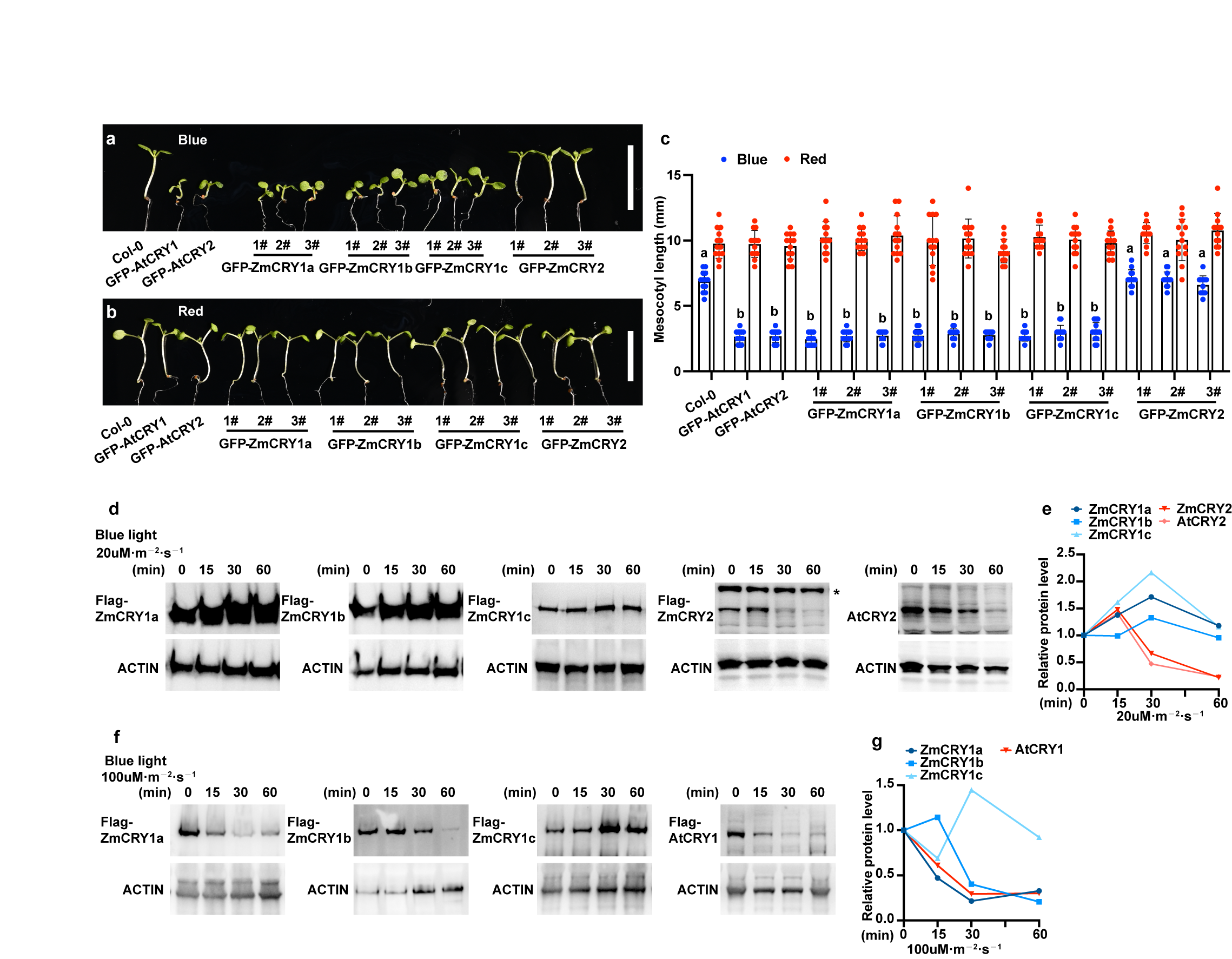

Supplementary Fig. 3 ZmCRY1s have conserved function to AtCRYs as blue light photoreceptor. related to Fig. 1.

**(a and b)**, Representative photographs of 8-day old of indicated genotypes grown under blue light (a, 20 μmol·m^−2^·s^−1^) and red light (b, 20 μmol·m^−2^·s^−1^). **(c)**, Hypocotyl lengths of the indicated genotypes shown in A and B. Error bars represent the s.d. (*n* = 13). Lowercase letters indicate statistically significant differences, no significant differences between genotypes under red light, as determined by one-way ANOVA with Tukey’s multiple comparisons test (*P* < 0.05). **(d to g)**, Immunoblot showing the ZmCRYs protein level in response to different intensities of blue light in their overexpression lines in *Arabidopsis*. 7-day old LD grown seedlings were pretreated with 48 h dark, then moved to medium blue light (20 μmol·m^−2^·s^−1^) (d and e) or high blue light (100 μmol·m^−2^·s^−1^) (f and g) for 1 hours’ time course. *: non-specific bond. ACTIN was used as a loading control. quantification of the relative CRY/ACTIN protein levels was show in e (for d) and g (for f). Dark levels were set as 1.

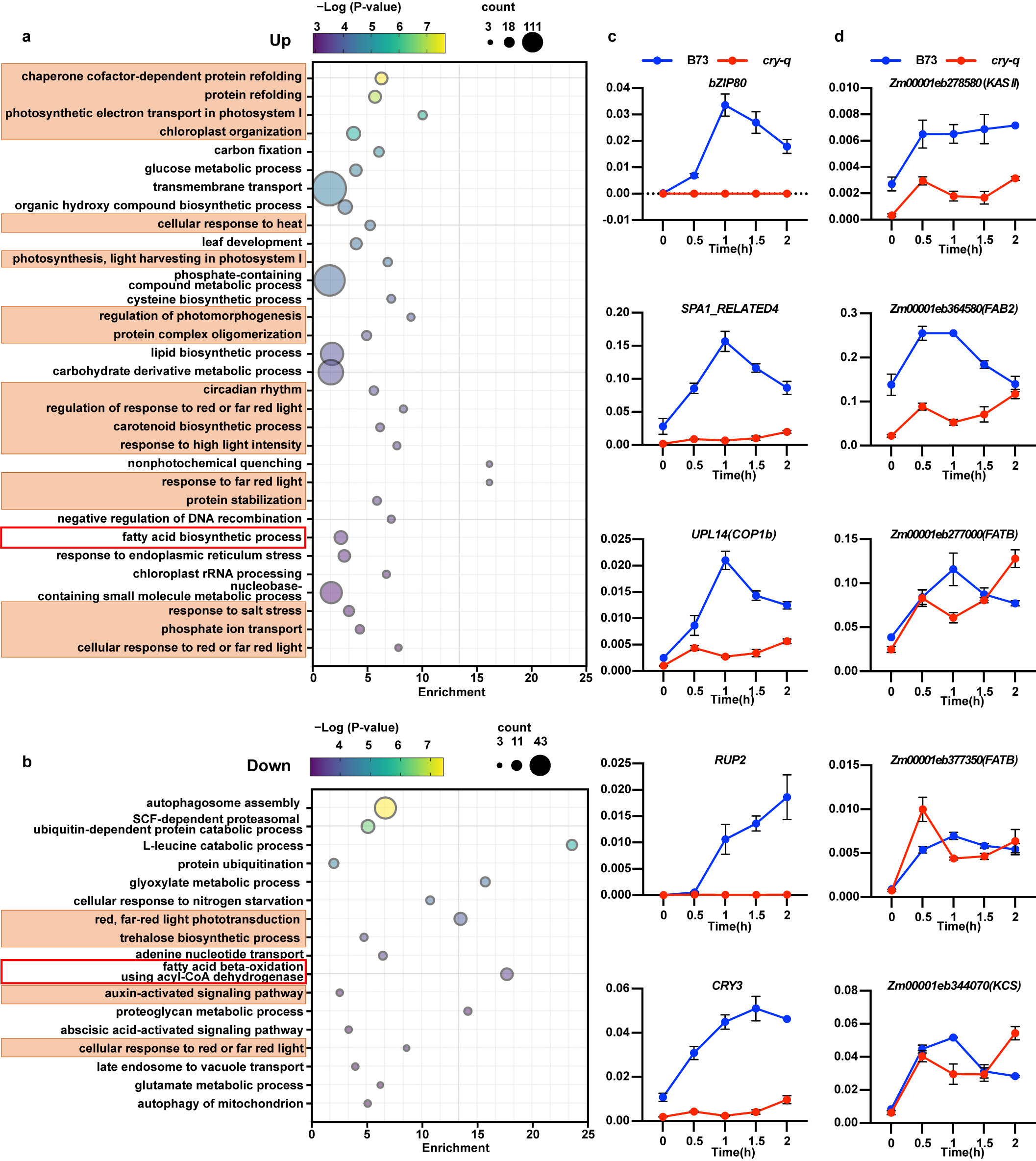

Supplementary Fig. 4 GO enrichment analysis of blue light regulated gene expression, related to Fig. 2 and Table 2.

**(a and b)**, GO enrichment items of up-regulated (a) and down-regulated (b) genes in blue light signaling identified in maize. The same blue light regulated GO items in *Arabidopsis* are highlighted. **(c and d)**, RT-qPCR validation the already known blue light signaling pathway regulators (c) and the up-regulated fatty acid biosynthesis enzymes (d) detected in the transcriptome. 7-day LD grown B73 and *Zmcry-q*, pretreated with 48h dark, then moved to blue light (20 μmol·m^−2^·s^−1^) for 2 hours’ time course. *ZmUBQ* served as internal control. Error bars, s.d. of three biological replicates.

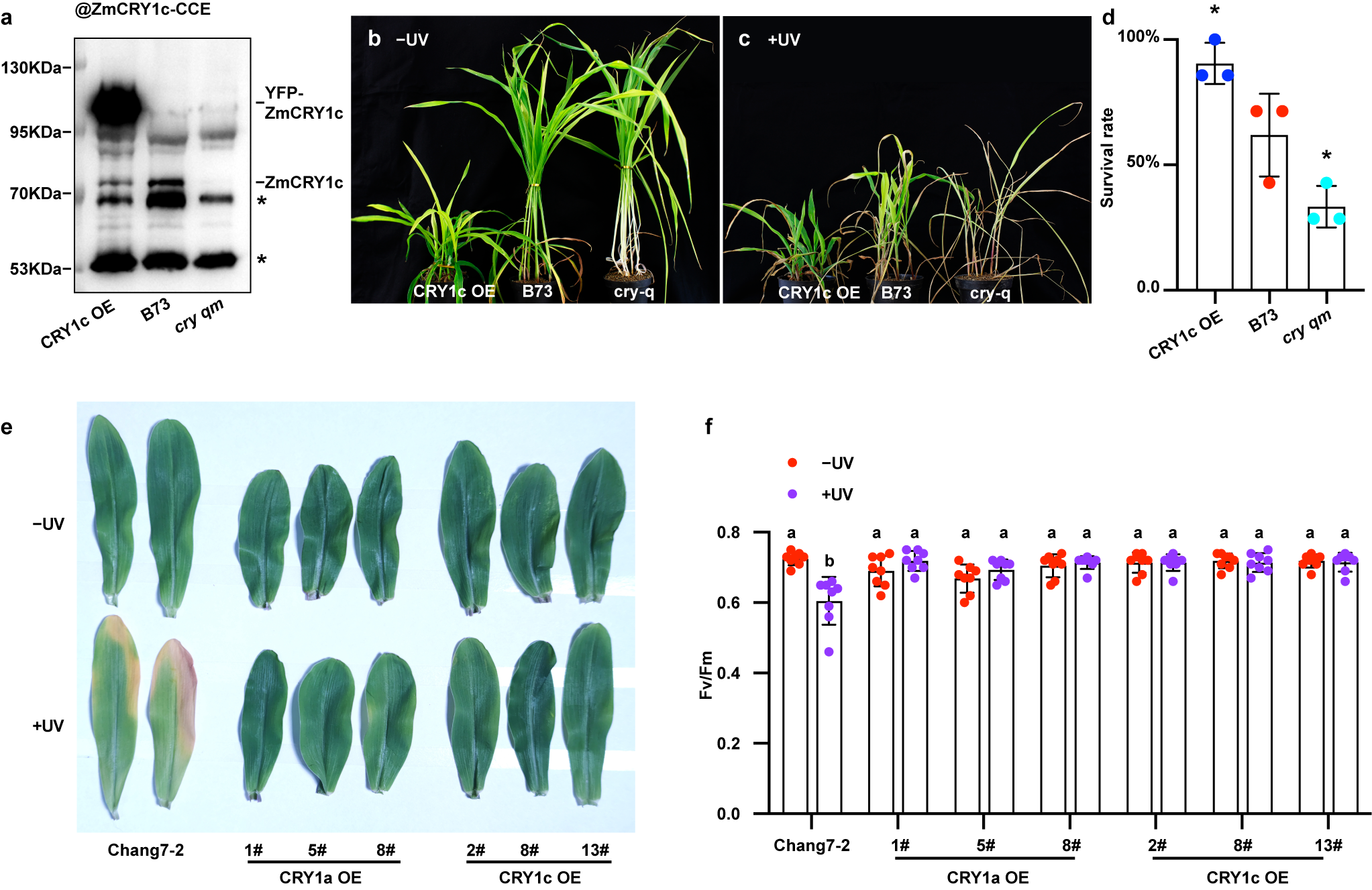

Supplementary Fig. 5 ZmCRYs are involved in UV-B stress tolerance in maize, related to Fig. 3.

**(a)**, Immunoblot showing the ZmCRY1c protein level in the indicated genotypes used in Fig.1E to H. *: non-specific bond. (**b to d)**, Analysis of ZmCRYs-mediated blue light UV-B stress tolerance in B73 background. Plants of the indicated genotypes were grown under LD in blue light (20 μmol·m^−2^·s^−1^) for 10 days and irradiated with (+UV) or without (−UV) broadband UV-B (5 μmol·m^-2^·s^-1^) for 6 hours on day 11 and allowed to recover for 2 days in blue light. **(d)**, Survival rate of indicated genotypes after UV-B stress. The asterisks indicate a significant difference from B73 based on one-way ANOVA with Dunnett’s multiple comparisons test (*P<0.05). **(e and f**), CRY1c OE show increased tolerance to UV-B stress. **(e)**, Analysis the UV-B stress tolerance of CRY1 OE in Chang7-2 background. Plants of the indicated genotypes were grown in soil under long-day (16/8) conditions in blue light (20 μmol·m^−2^·s^−1^) for 10 days and irradiated with (+UV) or without (−UV) broadband UV-B (5 μmol·m^-2^·s^-1^) for 6 hours on day 11 and allowed to recover for 2 days in blue light. **(f)**, Measurement of PSII maximal quantum yield (Fv/Fm) of the first leaves of the indicated genotypes (e). Fv/Fm was measured and quantified with an imaging fluorometer. Error bars represent standard deviation. Lowercase letters indicate statistically significant differences, as determined by two-way ANOVA with Tukey’s multiple comparisons test (P < 0.05).

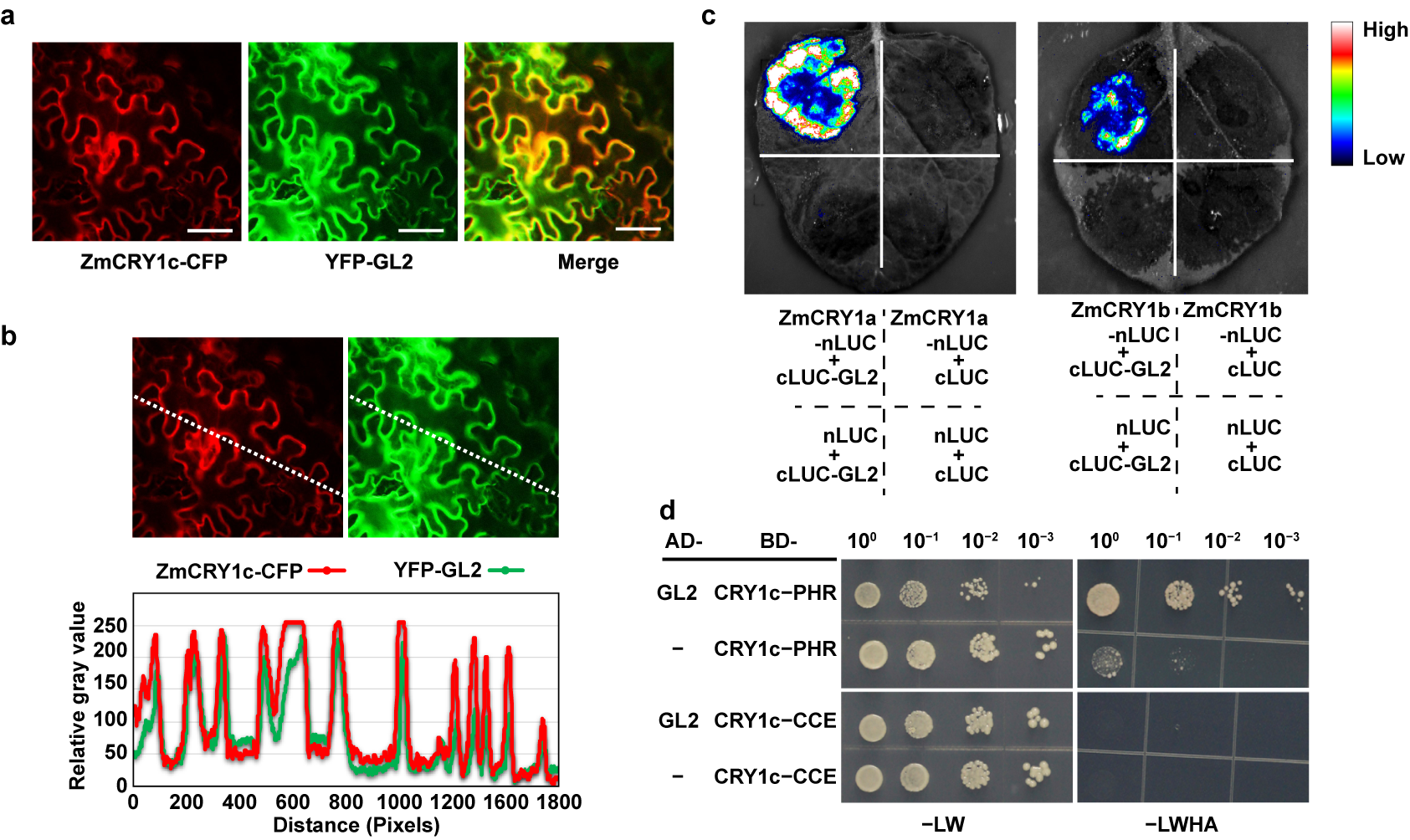

Supplementary Fig. 6 ZmCRY1s interact with GL2, related to Fig. 4.

(**a**), Co-localization of ZmCRY1c and GL2 in *N. benthamiana* leaves. Leaf epidermal cells of *N. benthamiana* were co-transformed to express the fusion proteins as indicated. In C, Scale bar = 50 μm. **(b)**, Gray-value analysis of fluorescent signals along the dashed diagonal line in the images at left. (**c**), BiLC assay showing *in vivo* protein interactions between ZmCRY1a and/or ZmCRY1b with GL2. (**d**), Histidine auxotrophy assays showing ZmCRY1c interacts with GL2 by its N-terminal PHR domain. Yeast cells were grown at –LW and –LWHA mediums.

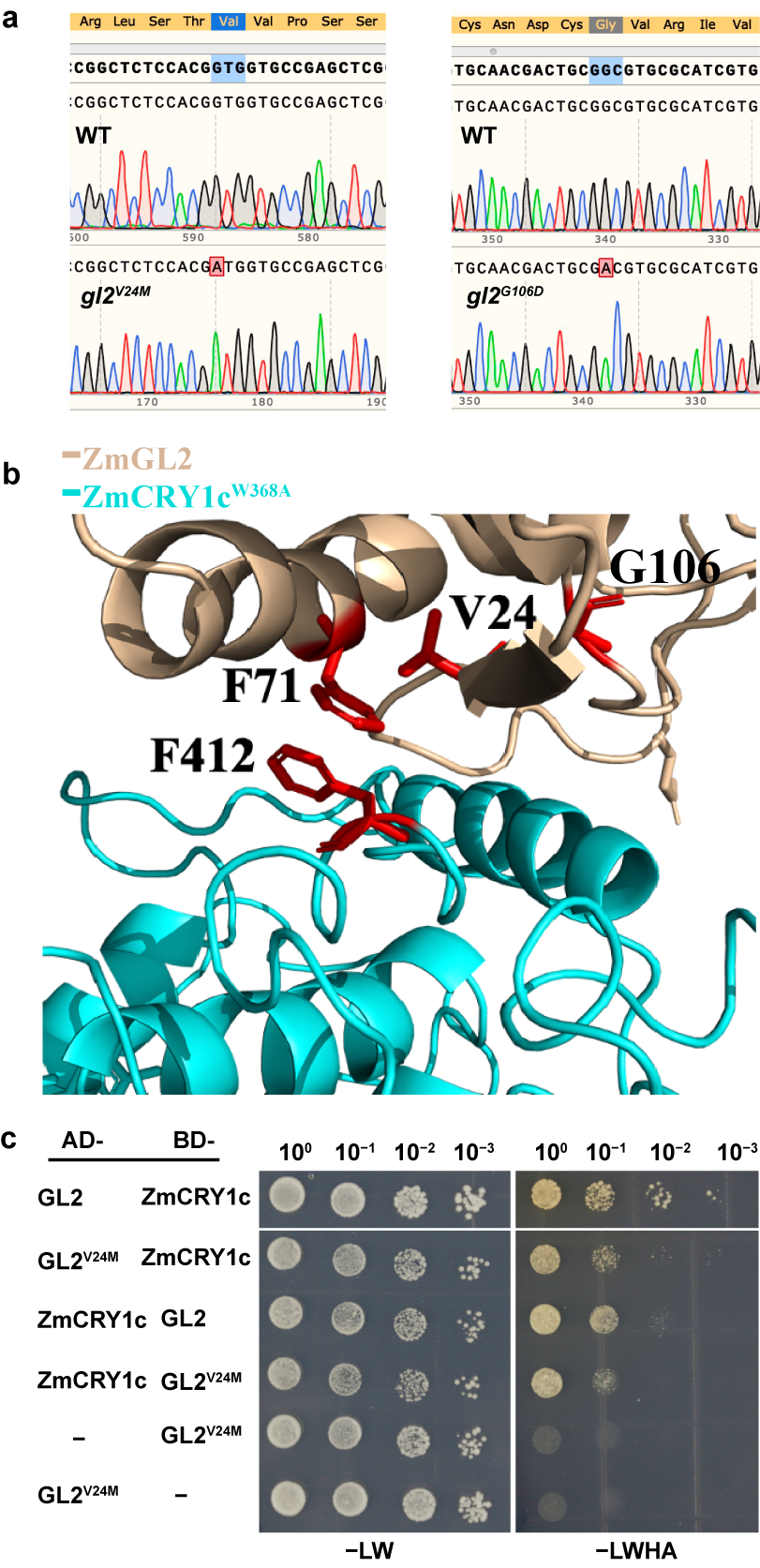

Supplementary Fig. 7 V24M and G106D of GL2 affect its’ interaction with ZmCRY1c, related to Fig. 5.

(**a**), Sanger sequencing chromatograms of the site mutation lines V24M and G106D of GL2 used in Fig.5. mutated bases are marked. (**b**), The tertiary structure of the interaction interfaces of ZmCRY1c and GL2 complex. The relative amino acid involved here are labeled in red color. (**c**), Y2H histidine auxotrophy assays showing the site mutation V24M of GL2 slightly affect its’ interaction with ZmCRY1c. Yeast cells were grown at –LWHA medium under blue light (30 μmol·m^–2^·s^–1^).

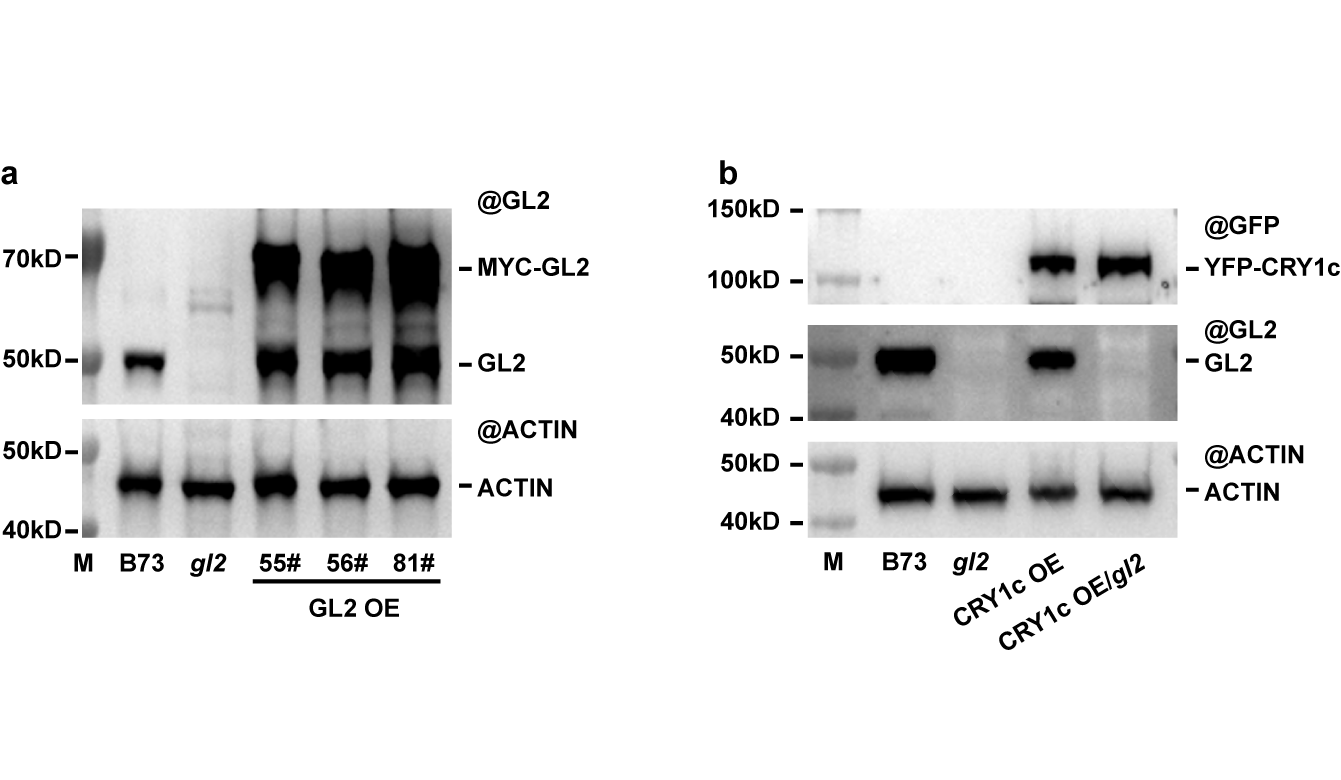

Supplementary Fig. 8 Identification of the MYC-GL2 and CRY1c OE/gl2 overexpression lines, related to Fig. 6.

(**a**), Immunoblots showing the MYC-GL2 protein level in their overexpression transgenic lines in B73 background as indicated. (**b**), Immunoblots showing the YFP-ZmCRY1c protein level in the indicated genotypes. ACTIN serves as the loading control.
